# Molecular basis of small-molecule binding to α-synuclein

**DOI:** 10.1101/2021.01.22.426549

**Authors:** Paul Robustelli, Alain Ibanez-de-Opakua, Cecily Campbell-Bezat, Fabrizio Giordanetto, Stefan Becker, Markus Zweckstetter, Albert C. Pan, David E. Shaw

**Affiliations:** D. E. Shaw Research, New York, NY 10036, USA; German Center for Neurodegenerative Diseases (DZNE), 37077 Göttingen, Germany; Max Planck Institute for Biophysical Chemistry, 37077 Göttingen, Germany; DFG Research Center for Nanoscale Microscopy and Molecular Physiology of the Brain (CNMPB), 37073 Göttingen, Germany; Department of Biochemistry and Molecular Biophysics, Columbia University, New York, NY 10032, USA

## Abstract

Intrinsically disordered proteins (IDPs) are implicated in many human diseases. They have generally not been amenable to conventional structure-based drug design, however, because their intrinsic conformational variability has precluded an atomic-level understanding of their binding to small molecules. Here we present long-timescale, atomic-level molecular dynamics (MD) simulations of monomeric α-synuclein (an IDP whose aggregation is associated with Parkinson’s disease) binding the small-molecule drug fasudil in which the observed protein-ligand interactions were found to be in good agreement with previously reported NMR chemical shift data. In our simulations, fasudil, when bound, favored certain charge-charge and π-stacking interactions near the C terminus of α-synuclein, but tended not to form these interactions simultaneously, rather breaking one of these interactions and forming another nearby (a mechanism we term *dynamic shuttling*). Further simulations with small molecules chosen to modify these interactions yielded binding affinities and key structural features of binding consistent with subsequent NMR experiments, suggesting the potential for MD-based strategies to facilitate the rational design of small molecules that bind with disordered proteins.

## Introduction

Intrinsically disordered proteins (IDPs), which lack a fixed three-dimensional structure under native, functional conditions, play important roles in a large number of biological pathways.^1–6^ IDPs and proteins with large disordered regions represent approximately 40% of the protein-coding human genome,^1,7^ and are also crucial components of biomolecular condensates, which have been increasingly recognized to be important regulators of cellular processes.^8^ IDPs are implicated in many human diseases, such as cancer, cardiovascular disease, diabetes, and neurodegeneration, and represent a large pool of potential drug targets.^9–15^ Drugging IDPs, however, has proven difficult due to their highly conformationally dynamic nature and the challenges associated with experimentally characterizing their conformational ensembles at atomic resolution.^16–22^ Because IDPs generally cannot be meaningfully represented by a single dominant conformation, or even a small number of substantially populated conformations, they are generally not suitable targets for conventional structure-based drug design methods, in which small molecules are designed to optimize interactions with a particular binding pocket in a folded protein.^10–13,23^

The aggregation of the IDP α-synuclein (α-syn) into oligomers and amyloid fibrils may play an important role in the etiology of Parkinson’s disease,^21,24^ and a potential therapeutic strategy for Parkinson’s disease is the stabilization of α-syn in its soluble monomeric form. Recently, the small molecule fasudil has been shown to interact with monomeric α-syn and delay its aggregation. Solution nuclear magnetic resonance (NMR) experiments demonstrated that this interaction is primarily localized to a specific cluster of protein residues at the C terminus.^25^ In contrast to a “lock-and-key” picture of protein-ligand recognition, in which a ligand binds to a stable and well-defined binding site, this fairly specific interaction between the ligand and the protein occurs while the protein remains highly dynamic and disordered. A detailed molecular picture of how fasudil binds to α-syn could shed light on how fasudil recognizes a specific region of disordered α-syn monomer, and provide a basis for the rational design of molecules that bind more strongly.

Atomic-level molecular dynamics (MD) simulations have been a valuable tool for complementing experimental measurements of disordered proteins and providing detailed descriptions of their conformational ensembles.^26–31^ Recent improvements to molecular mechanics potential energy functions, or “force fields,” have dramatically improved the accuracy of MD simulations of disordered proteins as assessed by their agreement with a wide variety of experimental measurements.^32–41^ MD simulations with these improved force fields have shown promise for describing molecular recognition mechanisms of IDPs in scenarios such as folding-upon-binding,^42^ dimerization,^43^ and the formation of higher-order molecular assemblies.^44,45^ MD simulations may also provide a promising approach for describing the binding of small molecules to disordered proteins in atomistic detail and for investigating the driving forces of these interactions.^11,23,46–53^

Here we report long-timescale MD simulations of fasudil binding to α-syn. The probability of observing contacts between fasudil and α-syn correlated remarkably well in our simulations with the magnitude of NMR chemical shift perturbations measured from α-syn–fasudil titrations. This correlation suggests that these simulations provide a highly accurate description of the molecular interactions that give rise to the preferential binding of fasudil to the C-terminal region of α-syn, which we found to be driven mainly by combinations of aromatic stacking and charge-charge interactions. The simulations provide an atomically detailed picture of the protein-ligand binding ensemble—in which α-syn remained largely disordered while fasudil transitioned between several distinct binding modes—and illustrate how a set of weak intermolecular interactions can lead to a small molecule interacting with a specific part of a protein that remains highly dynamic upon binding. We observed that although fasudil had a preference for a set of specific protein-ligand interactions, multiple such interactions rarely formed simultaneously. Instead, binding occurred through what we refer to as a *dynamic shuttling* mechanism, in which one of these preferred interactions broke before another formed nearby.

To better understand small-molecule features that confer affinity and specificity and to prospectively test the accuracy of our MD models, we simulated α-syn with a library of 49 small molecules that were selected to probe the simulated binding interactions and modify protein-ligand affinity. Subsequent NMR measurements of a subset of these small molecules showed that their relative binding affinities were in line with our computational predictions, and also provided support for key structural features of the simulated binding interactions, such as the populations of intermolecular hydrogen bonds and of aromatic stacking interactions. These observations illustrate that MD simulations can be a valuable tool for describing the binding of small molecules to disordered proteins, and suggest potential strategies for the rational design of molecules that bind disordered protein sequences with higher affinity and greater specificity.

## Results and Discussion

We performed a 1.5-ms MD simulation of α-syn and fasudil using the a99SB-*disp* force field^33^ and the generalized amber force field (GAFF) for fasudil.^54,55^ The probability of observing contacts between fasudil and each residue of α-syn is shown in Figure 1A. We also measured backbone ^15^N and ^1^HN NMR chemical shift perturbations (CSPs) of α-syn in the presence and absence of 2.7 mM fasudil. CSPs are sensitive to changes in the local chemical environment of each backbone amide bond, and are thus sensitive probes of protein-ligand interactions. The magnitude of the NMR CSPs is shown in Figure 1B. In agreement with previous measurements,^25^ we observed CSPs throughout the entire sequence of α-syn, with the largest-magnitude CSPs observed in the C-terminal residues 121–140.

**Figure 1.**
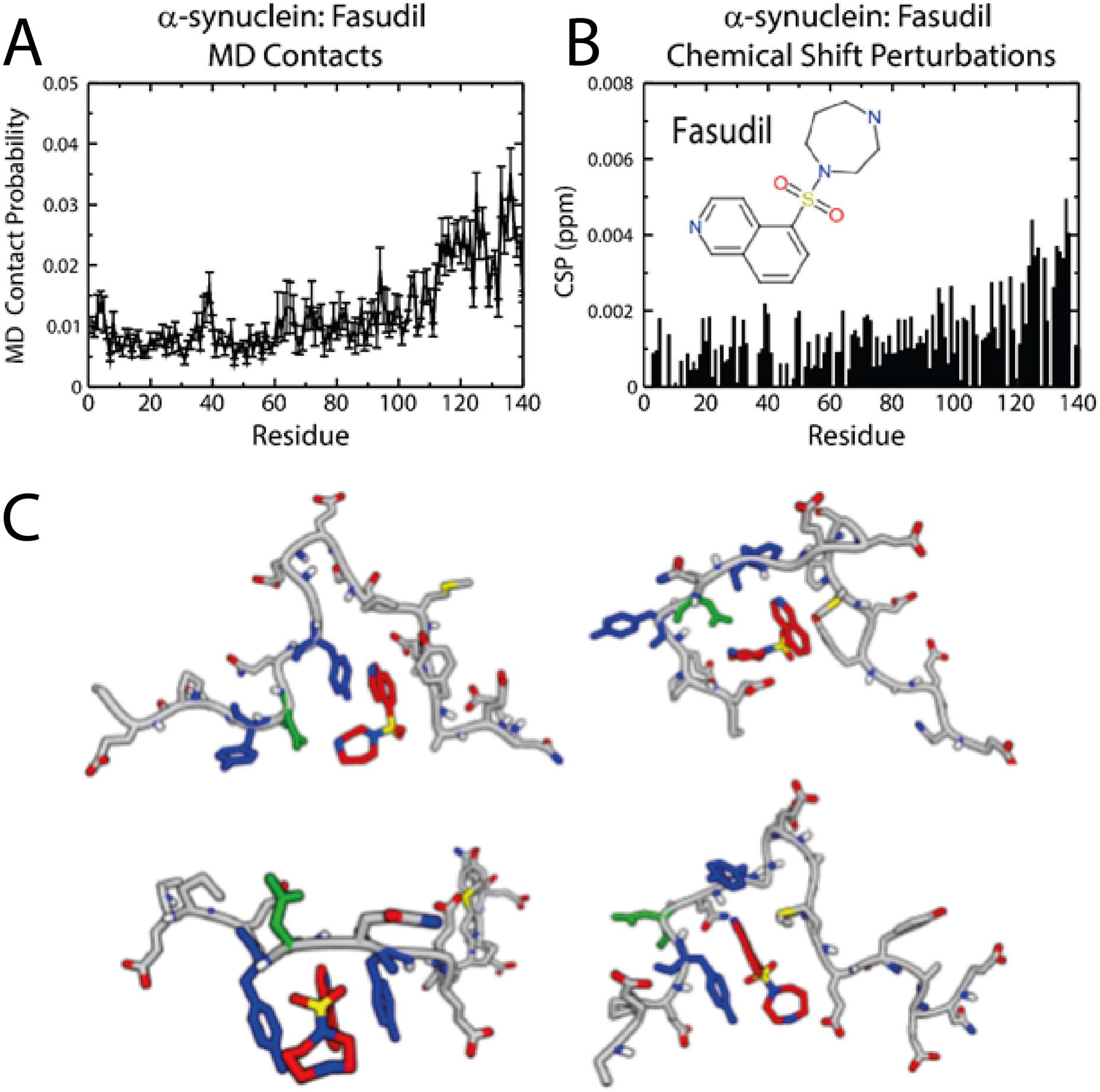
The dynamic binding mechanism of fasudil with α-synuclein observed in MD simulations is consistent with NMR chemical shift perturbation experiments. A) Contact probabilities between each residue of α-synuclein and fasudil observed in an unbiased 1.5-ms MD simulation run with the a99SB-*disp* force field. A contact is assigned to all MD frames when the minimum distance between any atom of fasudil and any heavy atom of a protein side-chain residue is <6 Å. B) NMR chemical shift perturbations of α-synuclein measured in the presence of 2.7-mM fasudil. C) Snapshots of binding modes of fasudil (red carbons) with α-syn-C-term, illustrating the conformational diversity of the bound ensemble. The residues with the highest probability of interacting with fasudil in the bound ensemble are colored blue (Y133, Y136) and green (D135).

The probability of observing contacts between fasudil and α-syn in the simulation and the magnitude of NMR CSPs measured for each residue are in excellent agreement (Figure S1, Pearson correlation coefficient, *r* = 0.67). Consistent with the experimental CSPs, in simulation we observed that fasudil interacts somewhat weakly with the entire α-syn sequence, but has a higher affinity for residues 121–140. The relatively small magnitude of the CSPs observed suggests that the underlying conformational ensemble of α-syn is not substantially altered in the presence of fasudil, and thus that α-syn remains disordered while interacting with fasudil. In simulation, we also found no large-scale differences between the conformational ensembles of the bound and unbound states of α-syn (Figures S2 and S5)–nor between the conformational ensembles of bound and unbound states of fasudil (Figure S2)—and that the bound ensemble is not dominated by a single conformation or a small number of substantially populated conformations (Figure S3). These results illustrate that the recently developed a99SB-*disp* force field,^33^ which provides improved descriptions of disordered proteins, is capable of identifying the binding sites of a small molecule in the context of an entire disordered protein sequence, and suggest that our simulation provides a meaningful model of the interactions between α-syn and fasudil.

In order to obtain a better understanding of the bound protein-ligand ensemble and the intermolecular interactions that confer specificity of fasudil to the C terminus of α-syn, we performed additional simulations of fasudil and a truncated α-syn construct containing only the region of α-syn to which fasudil preferentially binds (residues 121–140), which we will refer to as *α-syn-C-term*. Using this construct enabled more efficient simulation (given the smaller sizes of the protein and water box), which allowed us to obtain better statistics on the populations of dominant intermolecular interactions and the distributions of the α-syn conformations in bound states. Importantly, simulations with the reduced construct produced a similar contact probability with fasudil when compared to full-length α-syn (Figure S4). Simulations of full-length α-syn with fasudil also did not appear to involve long-range protein contacts that influenced small-molecule binding (Figure S5), implying that the simulation of full-length α-syn may not be required to model how fasudil binds to the C-terminal region and further justifying the use of a smaller protein construct. Similarly to simulations of full length α-syn, simulations of fasudil with α-syn-C-term did not give rise to substantially populated long-range protein contacts (Figure S5) or display a substantial difference between the conformational ensembles of bound and unbound states (Figure S6).

We found that, although bound protein-ligand conformations (i.e., those in which there was at least one contact between the protein and the ligand) exhibited a heterogeneity of binding modes (Figure 1C) instead of a single, stable protein-ligand complex (or a small number of them), we did observe a preference for specific charge-charge and aromatic π-stacking protein-ligand interactions near the C-terminus (Figure 2). We calculated the probability of observing hydrophobic contacts, charge-charge interactions, aromatic π-stacking interactions, and hydrogen bonds between fasudil and each residue of α-syn-C-term in bound conformations (Figure 2A) and found that the relative populations of protein-ligand interactions varied among similar amino acids in the α-syn sequence. That is, aromatic side chains did not all have the same propensity to form hydrophobic contacts and aromatic π-stacking interactions, and not all negatively charged side chains had the same propensity to form charge-charge interactions with fasudil.

**Figure 2.**
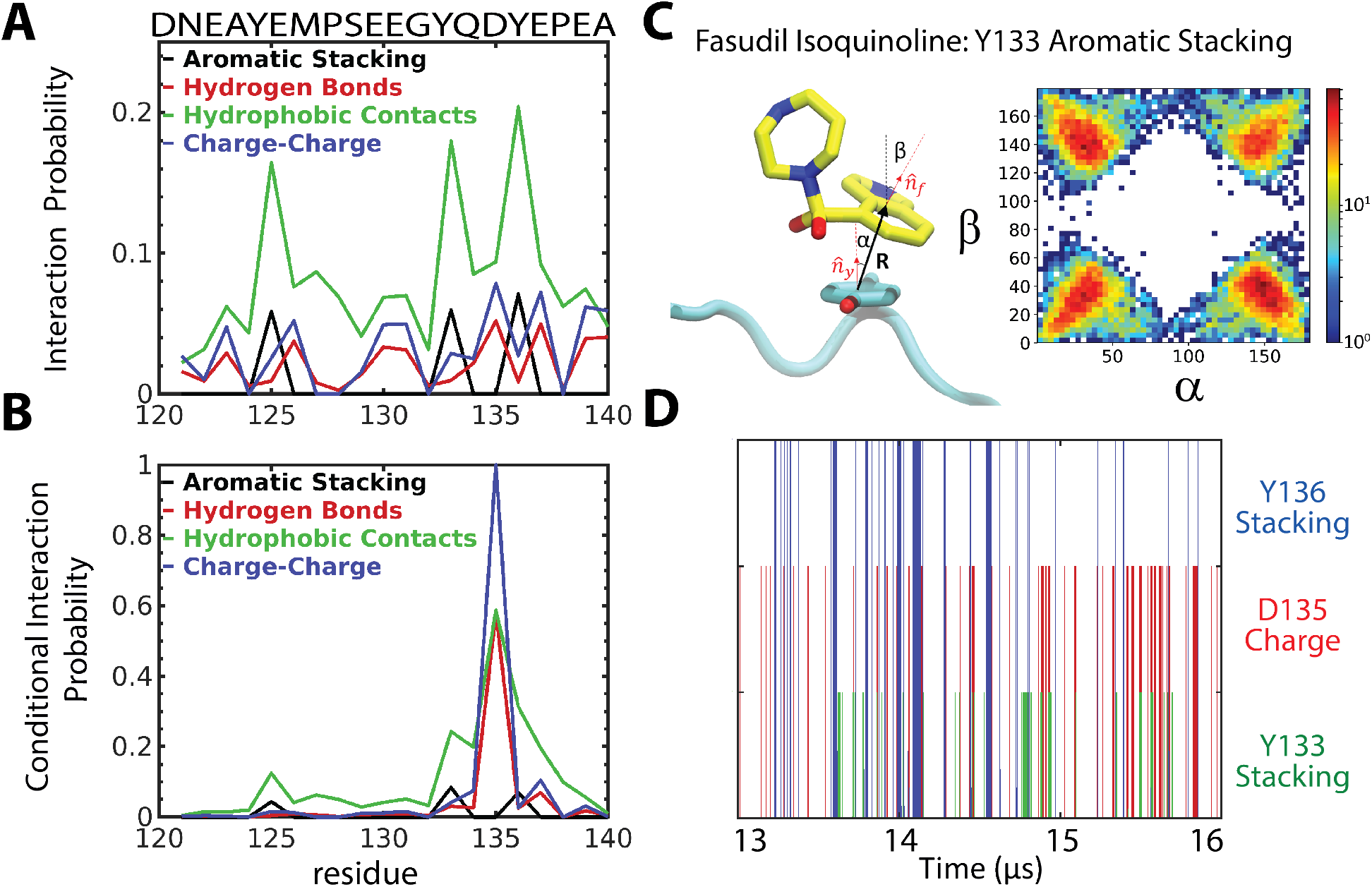
When in contact with α-synuclein, fasudil dynamically shuttled between different binding modes with different interactions; multiple specific interactions were rare in any given binding mode. A) The probability of observing interactions between fasudil and α-syn-C-term categorized by type of interaction in the bound ensemble. We note that a given residue can only form certain types of interactions. B) The conditional interaction probability of observing a specific interaction between α-synuclein and fasudil in the bound ensemble, given that a charge-charge interaction had formed between D135 and fasudil. C) An illustration of the stacking orientation between fasudil’s isoquinoline ring and the Y133 side chain. **R** is the distance vector between the centers of mass of the six aromatic carbons of Y133 and ten aromatic atoms of the isoquinoline ring on fasudil. The distributions are normalized and shown on a logarithmic scale. D) A time series of a representative portion of the unbiased MD trajectory of α-synuclein with fasudil showing the formation of different interactions. The presence of a line indicates the formation of a particular interaction. Trajectory frames were sampled every 180 ps.

Excluding hydrophobic contacts, which are relatively non-specific, the most likely interactions were a charge-charge contact between the positively charged amine of fasudil’s azepane ring and the side chains of D135 and E137, and aromatic π-stacking interactions between fasudil’s isoquinoline ring and the side chains of Y133 and Y136. In particular, D135 was approximately twice as likely to form a charge-charge interaction with fasudil as was E126 (Figure 2A).

Examination of the bound-state ensemble suggests that increased affinity between fasudil and specific residues in the α-syn-C-term sequence cannot be explained by the simultaneous formation of multiple specific interactions within individual conformations of the bound ensemble. If we consider only conformations where fasudil had a charge-charge interaction with D135, for example (the most likely charge-charge interaction between the protein and the ligand (Figure 2A)), very few of those conformations also made an additional specific protein-ligand interaction (Figure 2B). Only three interactions—aromatic stacking interactions with Y133 and Y136, and charge-charge interactions with E137—were formed in >5% of MD frames in which fasudil also formed charge interactions with D135. These observations about the lack of multiple specific protein-ligand interactions in the bound ensemble are also corroborated by a mutual information analysis: Calculating the mutual information between different protein-ligand interactions shows very little coupling between different interactions (Figure S7). Instead, we observed that the formation of a single intermolecular contact often spatially localized and oriented fasudil such that additional intermolecular interactions, although not forming simultaneously, became accessible later in time with relatively small ligand displacements.

To better understand the nature of the coupling among interactions, we sought to determine the extent to which the formation of specific intermolecular interactions could geometrically orient fasudil relative to α-syn, even as α-syn remained highly dynamic. We thus examined the distribution of π-stacking orientations^56^ of the isoquinoline ring of fasudil with side chains of Y125, Y133, and Y136 to determine how the identity of neighboring residues influenced the distribution of orientations of these interactions. For all conformations in which the ring centers of the fasudil isoquinoline group and a tyrosine phenol group were within 5 Å, we calculated the orientations of the normal vectors of each ring plane relative to a vector connecting the two ring centers (Figure 2C, left panel). If fasudil were interacting with an isolated tyrosine with no preferred geometric orientation, one would expect the four quadrants of a graph representing the distribution of the orientations of the ring plane normal vectors to be equally populated. That is, the isoquinoline ring would have an equal probability of stacking above or below the tyrosine phenol ring, and would have an equal probability of facing upwards or downwards in either position. We observed that, although the distribution of π-stacking orientations between fasudil and Y125 was indeed relatively symmetric, there was a greater asymmetry in the π-stacking orientations between fasudil and the C-terminal tyrosine residues, Y133 and Y136 (Figures 2C and S8; Table S1).

The distribution of π-stacking orientations observed between fasudil and Y133 is shown in Figure 2C (right panel). We observed that the most populated stacking orientation (α > 90, β < 90, quadrant 4 in Table S1) occurred 150% more frequently than did the least populated orientation (α > 90, β > 90, quadrant 2 in Table S1). We also examined the distribution of π-stacking orientations for just the conformations in which fasudil formed a charge-charge contact with D135 (Table S1). In these conformations the asymmetry of π-stacking orientations was even more pronounced: ~460% enrichment for (α > 90, β < 90) relative to (α < 90, β < 90) (Table S1). A similar effect—enrichments of 220% for (α > 90, β > 90) relative to (α > 90, β < 90) and 410% for (α < 90, β > 90) relative to (α > 90, β < 90)—was observed for Y136 and Y125, respectively, in the presence of D135 charge contacts. Although fasudil rarely formed both a π-stacking interaction with Y133 and a charge-charge contact with D135 simultaneously (<1% of frames in the bound ensemble contain both these interactions), the fact that the distribution of π-stacking orientations observed between fasudil and Y133 was asymmetric even in the absence of simultaneous charge-charge interactions with D135 suggests that the proximity and spatial orientation of Y133 and D135 may still confer a preferred set of orientations of fasudil relative to α-syn.

We illustrate this example of the *dynamic shuttling* of fasudil and α-syn between complementary intermolecular interactions in Figure 2D with a time series of the formation of different intermolecular interactions. Temporal correlations among these interactions persist for as long as 100 ns (Figure S8B). The probability, for example, of observing a π-stacking interaction between fasudil and Y136 after observing a charge-charge interaction between fasudil and D135 is greater than random for almost 100 ns, whereas the average lifetime of configurations with simultaneous interactions between fasudil and D135 and Y136 is only 570 ps. Although multiple interactions rarely formed simultaneously, fasudil transiently shuttles back and forth among favorable interactions, remaining localized to the same region of α-syn.

Having obtained a better understanding of the binding interactions between fasudil and α-syn, we proceeded to test our model prospectively by simulating the binding of additional small molecules and then subsequently measuring the CSPs of a subset of the compounds by NMR. We selected 49 commercially available compounds containing variations of the isoquinoline, sulfonyl, and azepane scaffold of fasudil (Figure S9). We expected that these variations would influence available binding modes and affinity to α-syn. We then conducted a computational screen, in which we performed 60-μs simulations of each molecule with the α-syn-C-term fragment, and calculated the simulated dissociation constant, *K*_*D*_, of each molecule (Table S3, Figure S9). The small molecule with the lowest *K*_*D*_ value (Ligand 47), two molecules with *K*_*D*_ values similar to fasudil (Ligands 2 and 5), and the ligand with the highest *K*_*D*_ value (Ligand 23) are shown in Figure 4A along with fasudil.

The simulation results for the screened compounds could in some cases be explained based on their structure and information we learned from our fasudil simulations. Charge-charge interactions between fasudil and α-syn were important binding interactions observed in our simulations, for example, and in our screening simulations, molecules that did not have a positive charge (such as Ligands 23 and 30) had higher *K*_*D*_ values, binding to α-syn-C-term ~2-fold more weakly compared to fasudil.

The simulation results of the strongest binder, Ligand 47, would have been more difficult to deduce from the structure of the compound alone. Ligand 47 differs from fasudil by a change in the substitution position of the sulfonyl group relative to the ring system, and the addition of an acetyl group to a slightly modified ring system, and it was not immediately clear how these features may have resulted in its ~2-fold higher affinity to α-syn-C-term. To obtain improved statistics for these interactions, the simulation of α-syn-C-term and Ligand 47 was extended to 200 μs. Examination of the interaction profile of Ligand 47 with α-syn-C-term (Figure 3A) shows that the largest differences in the bound ensembles were an increased propensity for π-stacking in Y136 (in 13.3% of bound conformations compared to 7.1% of bound conformations for Ligand 47 and fasudil, respectively) and the formation of hydrogen bonds between Ligand 47 and P128, Y136, and P138 (in 1%, 2.1%, and 1.9% of bound conformations, respectively) that were not appreciably populated in the bound fasudil ensemble.

**Figure 3.**
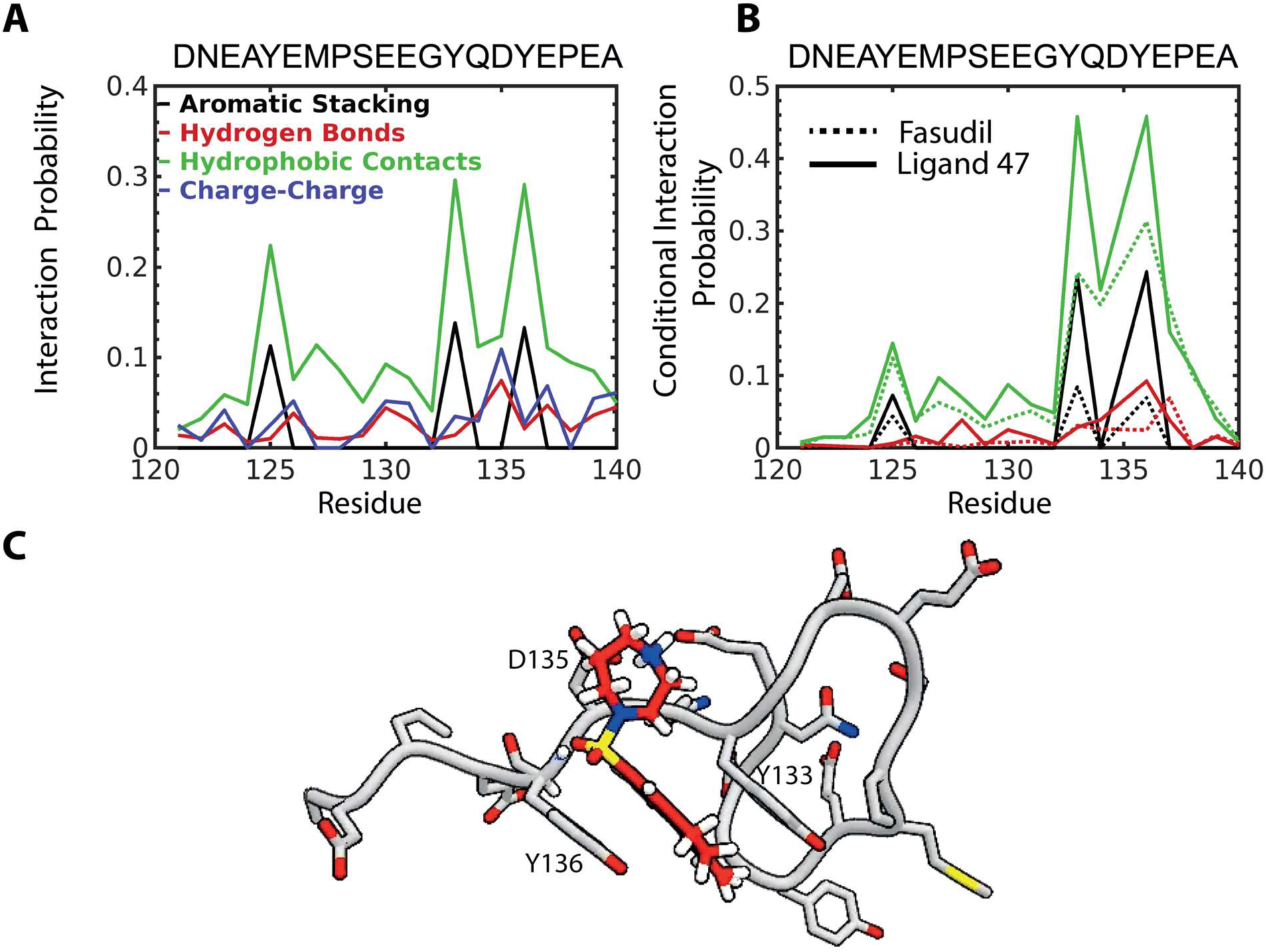
Predicted interactions of Ligand 47 with α-synuclein. A) Interaction probabilities for each residue 121–140 of α-synuclein and Ligand 47 observed in an unbiased MD simulation. B) The conditional interaction probability of observing interactions between α-synuclein and Ligand 47, and between α-synuclein and fasudil, for all conformations containing a charge-charge contact with D135. C) A representative structure of the most populated cluster of conformations of the Ligand 47 bound ensemble. In this conformation, Ligand 47 (red carbons) can simultaneously stack with Y136, form a hydrogen bond with the backbone amide of Y136, and form charge contacts with D135.

We also observed that the position of the sulfonyl group of Ligand 47 relative to the ring system, and the corresponding change in the orientation of the charged amine of the azepane ring, seemed to increase the probability of forming multiple intermolecular interactions with α-syn-C-term simultaneously. The average mutual information between different protein-ligand interactions was increased relative to fasudil (Figure S7), and we observed greater correlations between interaction pairs involving non-neighboring residues. In particular, we observed much stronger couplings among interaction pairs involving D135. In Figure 3B, we compare the conditional probabilities of observing additional intermolecular interactions for all conformations in which Ligand 47 forms a charge contact with D135 to the corresponding probabilities for fasudil. We observed that Ligand 47 had increased conditional probabilities of forming π-stacking interactions with Y136 (24.4% vs. 6.7%), hydrogen bond interactions with Y136 (9.2% vs. 2.5%), hydrophobic contacts with Y136 (45.8% vs. 31.3%), π-stacking interactions with Y133 (23.6% vs. 8.4%), and hydrophobic contacts with Y133 (45.8% vs 24.2%).

Examination of configurations in which Ligand 47 formed a charge interaction with D135 provides a possible explanation for these increases. The altered substitution position of the sulfonyl group oriented Ligand 47 such that when it formed a charge interaction with D135, there was a dramatic increase in the asymmetry of π-stacking orientations with Y136: With the D135 charge interaction, orientations with β < 90 were ~10-fold more populated than orientations with β > 90 (Table S1, Figure S8). This stacking orientation also oriented the sulfonyl oxygens such that they were able to form hydrogen bonds with the amide of Y136, and positioned the additional acetyl group such that it was able to form hydrophobic contacts with Y136. A representative bound conformation of Ligand 47 in its preferred stacking orientation and forming all 4 of these interactions is shown in Figure 3C.

We note that bound conformations in which Ligand 47 simultaneously formed a hydrogen bond with Y136, aromatic stacking interactions with Y136, and charge contacts with D135 constitute <0.9% of the bound ensemble. This suggests that conformations that can simultaneously form all of these interactions do not, by themselves, explain the increased affinity of Ligand 47 for α-syn relative to fasudil, as the small population of these states cannot account for the ~2-fold increase in affinity. Instead, the increased affinity is better explained in the context of the dynamic shuttling model, in which these interactions are part of a larger network of transient and weak interactions that localize Ligand 47 to these residues.

We next experimentally tested our computational predictions of relative binding affinities for a subset of screened compounds by measuring NMR CSPs of all α-syn residues in titrations of four ligands: Ligand 47, Ligand 23, Ligand 5, and Ligand 2 (Figure 4A). Due to solubility limitations of these molecules, it was not possible to saturate the observed CSPs and estimate *K*_*D*_s directly. We instead estimated the relative affinities of the molecules by considering the slope of the CSPs measured for Y125, Y133, and Y136 during each titration, with larger slopes indicating higher affinities. We found that by taking the average slope observed for these tyrosine residues as a proxy for affinity to the C-terminal region of α-syn, the relative affinities of the five ligands estimated from the ligand titrations are in line with the affinities predicted from our MD simulations.

**Figure 4.**
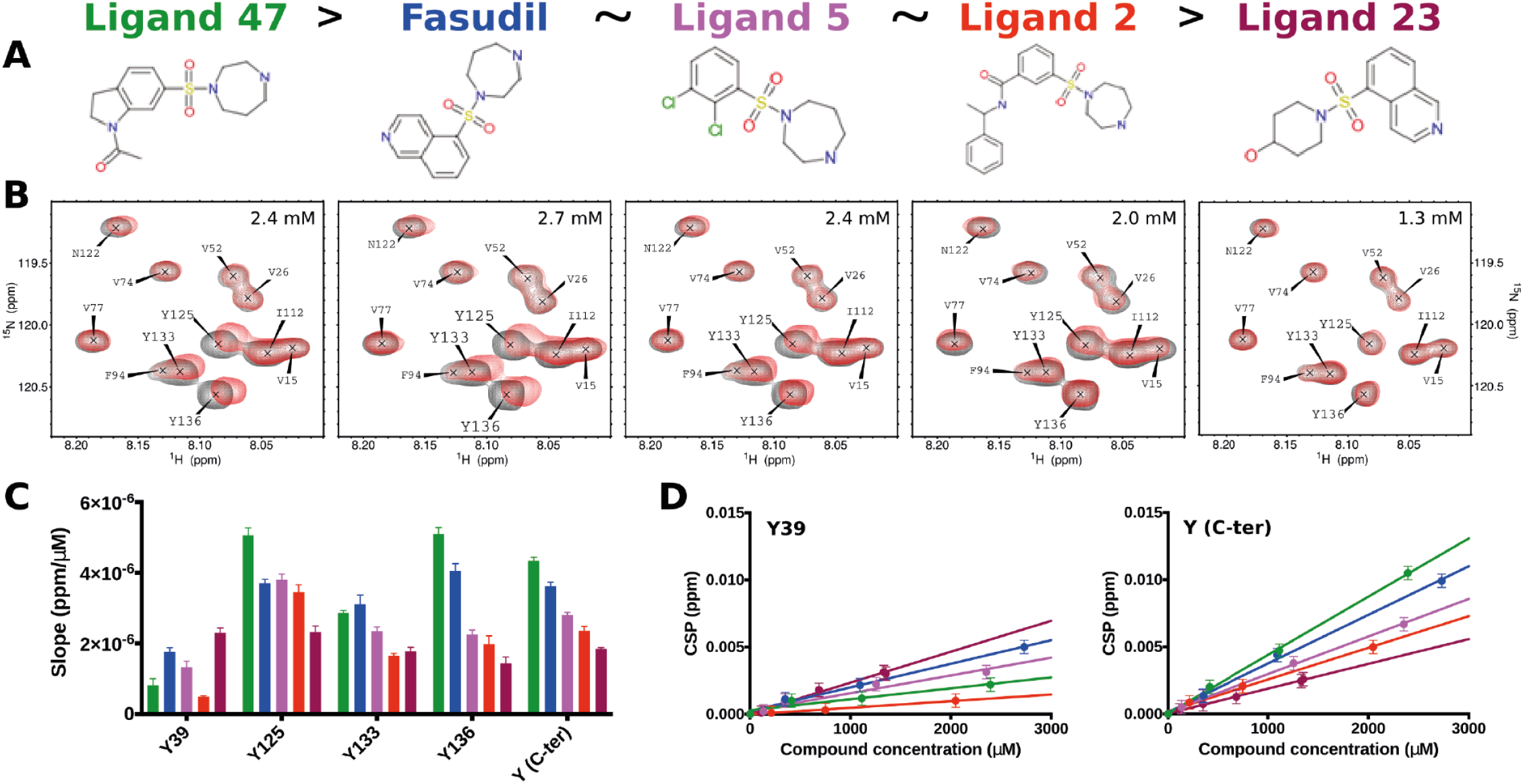
Predicted binding affinities of fasudil analogues with α-synuclein from simulation are in line with subsequently measured chemical shift perturbation titrations from NMR. A) Structures of fasudil and tested analogues, with Ligand 47 having the highest affinity for α-synuclein, and Ligand 23 the lowest. B) NMR chemical shift titration curves of the aromatic residues of the C-terminal region of α-synuclein with the five ligands depicted in A. C) Slope of titration curves for each tyrosine residue in α-synuclein, and the average of all tyrosine residues in the C-terminal region of α-synuclein (Y125, Y133, Y136). D) Titration curves of each compound for Y39, and for the average of all tyrosine residues in the C-terminal region of α-synuclein. Individual titration curves for Y125, Y133, and Y136, are shown in Figure S10.

Interestingly, we observe that Ligand 47 causes larger CSPs in Y136 and Y125 than does fasudil (Figure 4B, Figure S10). This is consistent with the interaction profiles observed from our MD simulations, in which Ligand 47 had a higher propensity to form hydrogen bonds and π-stacking interactions with Y136. The increased contact probability and stacking between Ligand 47 and Y125 are also consistent with Ligand 47’s higher affinity than fasudil for Y125 observed by NMR. To examine whether Ligand 47, like fasudil, had an increased propensity to interact with the C-terminal region of α-syn, we performed a 1.5-ms unbiased MD simulation of Ligand 47 with full-length α-syn. We observed in this simulation that Ligand 47, similarly to fasudil, had increased affinity for the C-terminal region of α-syn rather than exhibiting non-specific affinity for all residues, in agreement with observed NMR CSPs (r = 0.68, Figure S11, Table S2).

Lastly, the experimentally tested ligands showed varying levels of affinity for Y39 in the NMR CSP titration experiments, and differed from the relative affinities observed at the C-terminus (Figure 4). Y39 has been shown to be key residue for modulation of α-syn aggregation by small molecules^57,58^ as well as for α-syn dimerization in the oxidative environments persistent in disease states.^59^ Of the experimentally tested ligands, Ligand 23 showed the weakest affinity for the C-terminal region, something we also observed in the simulations, but had the strongest affinity for Y39 (Figure 4C). This prompted us to simulate the small molecules in our library with a fragment of α-syn from residues 29–49, in order to determine if our simulations would accurately capture the relative affinities between the tested compounds and Y39. We found that the simulations correctly identified Ligand 23 as the strongest binder to the α-syn 29–49 fragment among the experimentally tested ligands, but did not capture well the overall relative affinities of ligands at Y39 (Figure 4, Table S3).

We also examined the binding of Ligand 23 with full length α-syn. Although the affinity of Ligand 23 to Y39 in full-length α-syn is not the strongest relative to fasudil and Ligand 47, we did find that, in contrast to fasudil and Ligand 47, which heavily favored binding at the C-terminus relative to Y39, Ligand 23 had a slight preference for Y39 (Table S2), in agreement with experiment. These results suggest that our MD simulations may be capable of differentiating between the affinities of small molecules to tyrosine residues in different sequence contexts.

## Conclusion

We performed long-timescale MD simulations of fasudil binding to α-syn and found, in agreement with NMR chemical shift perturbations, that fasudil prefers to bind to the C-terminal region of α-syn—a preference driven mainly by a combination of aromatic stacking and charge-charge interactions. The simulations provide an atomically detailed picture of the protein-ligand binding ensemble, illustrating how a network of weak and transient intermolecular interactions can lead to specific binding of a small molecule to a protein that remains highly dynamic. The simulated binding mechanism observed here, in which the formation of specific intermolecular interactions orients ligands such that additional interactions are likely to form subsequently, provides a mechanistic explanation for how ligands can achieve sequence-specific binding with IDPs.

We note that the dynamic shuttling mechanism described here provides a local description of IDP-ligand binding to a given binding site. Dynamic binding mechanisms such as those characterized in this study could potentially underlie IDP-ligand binding events in which the presence of a small molecule either reduces the conformational space accessible to an IDP in its bound state (sometimes referred to as “conformational restriction,”^51^ “population-shifting,”^60^ or “entropic collapse”) or increases the conformational space accessible to an IDP in its bound state (sometimes referred to as “entropic expansion”^11,50^), and are not inherently associated with either thermodynamic binding scenario. The dynamic shuttling of a small molecule among heterogeneous binding modes could either stabilize a subset of conformations observed in the apo state of an IDP, or increase the sampling of additional conformations not substantially populated in the apo state.

In addition to confirming that simulations of fasudil and α-syn agreed retrospectively with previous NMR experiments, we prospectively tested the accuracy of our MD models by simulating α-syn with a library of 49 small molecules and testing the MD predictions with NMR measurements of a subset of these molecules. The experimentally determined relative binding affinities of these molecules were in line with our computational predictions, and the measured CSPs provided support for key structural features of the simulated binding interactions. These observations suggest potential strategies for the rational design of molecules that bind disordered protein sequences with higher affinity and greater specificity using insights from MD simulations.

## Acknowledgments

The authors thank Michael Eastwood and Stefano Piana for helpful discussions, Berkman Frank for editorial assistance, and Karin Giller and Melanie Wegstroth for excellent help with NMR sample preparation. M.Z. was supported by the advanced grant “787679 - LLPS-NMR” of the European Research Council.

## References

1. Van der Lee R, Buljan M, Lang B, Weatheritt RJ, Daughdrill GW, Dunker AK, Fuxreiter M, Gough J, Gsponer J, Jones DT, Kim PM, Kriwacki RW, Oldfield CJ, Pappu RV, Tompa P, Uversky VN, Wright PE, Babu MM. Classification of intrinsically disordered regions and proteins. Chem Rev. 2014 114(13):6589–6631.

2. Habchi J, Tompa P, Longhi S, Uversky VN. Introducing protein intrinsic disorder. Chem Rev. 2014 114(13):6561–6588.

3. Tompa P. Intrinsically unstructured proteins. Trends Biochem Sci. 2002 27(10):527–533.

4. Dunker AK, Cortese MS, Romero P, Iakoucheva LM, Uversky VN. Flexible nets. The roles of intrinsic disorder in protein interaction networks. FEBS J. 2005 272(20):5129–5148.

5. Dyson HJ, Wright PE. Intrinsically unstructured proteins and their functions. Nat Rev Mol Cell Biol. 2005 6(3):197–208.

6. Wright PE, Dyson HJ. Intrinsically disordered proteins in cellular signaling and regulation. Nat Rev Mol Cell Biol. 16(1): 18–29.

7. Sormanni P, Piovesan D, Heller GT, Bonomi M, Kukic P, Camilloni C, Fuxreiter M, Dosztanyi Z, Pappu RV, Babu MM, Longhi S, Tompa P, Dunker AK, Uversky VN, Tosatto SC, Vendruscolo M. Simultaneous quantification of protein order and disorder. Nat Chem Biol. 2017 13(4):339–342.

8. Banani SF, Lee HO, Hyman AA, Rosen MK. Biomolecular condensates: organizers of cellular biochemistry. Nat Rev Mol Cell Biol. 2017 18(5):285–298.

9. Babu MM, van der Lee R, de Groot NS, Gsponer J. Intrinsically disordered proteins: regulation and disease. Curr Opin Struct Biol. 2011 21(3):432–440.

10. Heller GT, Sormanni P, Vendruscolo M. Targeting disordered proteins with small molecules using entropy. Trends Biochem Sci. 2015 40(9):491–496.

11. Heller GT, Bonomi M, Vendruscolo M. Structural ensemble modulation upon small-molecule binding to disordered proteins. J Mol Biol. 2018 430(16):2288–2292.

12. Chen J, Liu X, Chen J. Targeting intrinsically disordered proteins through dynamic interactions. Biomolecules. 2020 10(5):743.

13. Fuertes G, Nevola L, Esteban-Martín S. Perspectives on drug discovery strategies based on IDPs. In “Intrinsically Disordered Proteins” (pp. 275–327). Academic Press, London, 2019.

14. Sadar MD. Discovery of drugs that directly target the intrinsically disordered region of the androgen receptor. Expert Opin Drug Discov. 2020 15(5):551–560.

15. De Mol E, Fenwick RB, Phang CT, Buzón V, Szulc E, de la Fuente A, Escobedo A, García J, Bertoncini CW, Estébanez-Perpiñá E, McEwan IJ, Riera A, Salvatella X. EPI-001, A compound active against castration-resistant prostate cancer, targets transactivation unit 5 of the androgen receptor. ACS Chem Biol. 2016 11(9):2499–2505.

16. Ruan H, Sun Q, Zhang W, Liu Y, Lai L. Targeting intrinsically disordered proteins at the edge of chaos. Drug Discov Today. 2019 24(1):217–227.

17. Metallo SJ. Intrinsically disordered proteins are potential drug targets. Curr Opin Chem Biol. 2010 14(4):481–488.

18. Iconaru LI, Ban D, Bharatham K, Ramanathan A, Zhang W, Shelat AA, Zuo J, Kriwacki RW. Discovery of small molecules that inhibit the disordered protein, p27(Kip1). Sci Rep. 2015 5:15686.

19. Bonomi M, Heller GT, Camilloni C, Vendruscolo M. Principles of protein structural ensemble determination. Curr Opin Struct Biol. 2017 42:106–116.

20. Jensen MR, Ruigrok RW, Blackledge M. Describing intrinsically disordered proteins at atomic resolution by NMR. Curr Opin Struct Biol. 2013 23(3):426–435.

21. Wu KP, Baum J. Detection of transient interchain interactions in the intrinsically disordered protein alpha-synuclein by NMR paramagnetic relaxation enhancement. J Am Chem Soc. 2010 132(16):5546–5547.

22. Sung YH, Eliezer D. Residual structure, backbone dynamics, and interactions within the synuclein family. J Mol Biol. 2007 372(3):689–707.

23. Jin F, Yu C, Lai L, Liu Z. Ligand clouds around protein clouds: a scenario of ligand binding with intrinsically disordered proteins. PLoS Comput Biol. 2013 9(10):e1003249.

24. Karpinar DP, Balija MB, Kügler S, Opazo F, Rezaei-Ghaleh N, Wender N, Kim HY, Taschenberger G, Falkenburger BH, Heise H, Kumar A, Riedel D, Fichtner L, Voigt A, Braus GH, Giller K, Becker S, Herzig A, Baldus M, Jäckle H, Eimer S, Schulz JB, Griesinger C, Zweckstetter M. Pre-fibrillar alpha-synuclein variants with impaired beta-structure increase neurotoxicity in Parkinson’s disease models. EMBO J. 2009 28(20):3256–3268.

25. Tatenhorst L, Eckermann K, Dambeck V, Fonseca-Ornelas L, Walle H, Lopes da Fonseca T, Koch JC, Becker S, Tönges L, Bähr M, Outeiro TF, Zweckstetter M, Lingor P. Fasudil attenuates aggregation of α-synuclein in models of Parkinson’s disease. Acta Neuropathol Commun. 2016 4:39.

26. Granata D, Baftizadeh F, Habchi J, Galvagnion C, De Simone A, Camilloni C, Laio A, Vendruscolo M. The inverted free energy landscape of an intrinsically disordered peptide by simulations and experiments. Sci Rep. 2015 5:15449.

27. Lindorff-Larsen K, Trbovic N, Maragakis P, Piana S, Shaw DE. Structure and dynamics of an unfolded protein examined by molecular dynamics simulation. J Am Chem Soc. 2012 134(8):3787–3791.

28. Tóth G, Gardai SJ, Zago W, Bertoncini CW, Cremades N, Roy SL, Tambe MA, Rochet JC, Galvagnion C, Skibinski G, Finkbeiner S, Bova M, Regnstrom K, Chiou SS, Johnston J, Callaway K, Anderson JP, Jobling MF, Buell AK, Yednock TA, Knowles TP, Vendruscolo M, Christodoulou J, Dobson CM, Schenk D, McConlogue L. Targeting the intrinsically disordered structural ensemble of α-synuclein by small molecules as a potential therapeutic strategy for Parkinson’s disease. PLoS One. 2014 9(2):e87133.

29. Salvi N, Abyzov A, Blackledge M. Solvent-dependent segmental dynamics in intrinsically disordered proteins. Sci Adv. 2019 5(6):eaax2348.

30. Salvi N, Abyzov A, Blackledge M. Analytical description of NMR relaxation highlights correlated dynamics in intrinsically disordered proteins. Angew Chem Int Ed Engl. 2017 56(45):14020–14024.

31. Robustelli P, Trbovic N, Friesner RA, Palmer AG 3rd. Conformational dynamics of the partially disordered yeast transcription factor GCN4. J Chem Theory Comput. 2013 9(11):10.1021/ct400654r.

32. Huang J, Rauscher S, Nawrocki G, Ran T, Feig M, de Groot BL, Grubmüller H, MacKerell AD Jr. CHARMM36m: an improved force field for folded and intrinsically disordered proteins. Nat Methods. 2017 14(1):71–73.

33. Robustelli P, Piana S, Shaw DE. Developing a molecular dynamics force field for both folded and disordered protein states. Proc Natl Acad Sci U S A. 2018 115(21):E4758–E4766.

34. Nerenberg PS, Jo B, So C, Tripathy A, Head-Gordon T. Optimizing solute-water van der Waals interactions to reproduce solvation free energies. J Phys Chem B. 2012 Apr 19;116(15):4524–4534.

35. Best RB, Zheng W, Mittal J. Balanced protein-water interactions improve properties of disordered proteins and non-specific protein association. J Chem Theory Comput. 2014 10(11):5113–5124.

36. Piana S, Donchev AG, Robustelli P, Shaw DE. Water dispersion interactions strongly influence simulated structural properties of disordered protein states. J Phys Chem B. 2015 119(16):5113–5123.

37. Piana S, Robustelli P, Tan D, Chen S, Shaw DE. Development of a force field for the simulation of single-chain proteins and protein-protein complexes. J Chem Theory Comput. 2020 16(4):2494–2507.

38. Tian C, Kasavajhala K, Belfon KAA, Raguette L, Huang H, Migues AN, Bickel J, Wang Y, Pincay J, Wu Q, Simmerling C. ff19SB: Amino-acid-specific protein backbone parameters trained against quantum mechanics energy surfaces in solution. J Chem Theory Comput. 2020 16(1):528–552.

39. Shabane PS, Izadi S, Onufriev AV. General purpose water model can improve atomistic simulations of intrinsically disordered proteins. J Chem Theory Comput. 2019 15(4):2620–2634.

40. Song D, Luo R, Chen HF. The IDP-specific force field ff14IDPSFF improves the conformer sampling of intrinsically disordered proteins. J Chem Inf Model. 2017 57(5):1166–1178.

41. Yu L, Li DW, Brüschweiler R. Balanced amino-acid-specific molecular dynamics force field for the realistic simulation of both folded and disordered proteins. J Chem Theory Comput. 2020 16(2):1311–1318.

42. Robustelli P, Piana S, Shaw DE. Mechanism of coupled folding-upon-binding of an intrinsically disordered protein. J Am Chem Soc. 2020 142(25):11092–11101.

43. Conicella AE, Dignon GL, Zerze GH, Schmidt HB, D’Ordine AM, Kim YC, Rohatgi R, Ayala YM, Mittal J, Fawzi NL. TDP-43 α-helical structure tunes liquid-liquid phase separation and function. Proc Natl Acad Sci U S A. 2020 117(11):5883–5894.

44. Paloni M, Bailly R, Ciandrini L, Barducci A. Unraveling molecular interactions in liquid-liquid phase separation of disordered proteins by atomistic simulations. J Phys Chem B. 2020 124(41):9009–9016.

45. Zheng W, Dignon GL, Jovic N, Xu X, Regy RM, Fawzi NL, Kim YC, Best RB, Mittal J. Molecular details of protein condensates probed by microsecond long atomistic simulations. J Phys Chem B. 2020 (in press).

46. Convertino M, Vitalis A, Caflisch A. Disordered binding of small molecules to Aβ(12-28). J Biol Chem. 2011 286(48):41578–41588.

47. Li G, Pomès R. Binding mechanism of inositol stereoisomers to monomers and aggregates of Aβ(16-22). J Phys Chem B. 2013 117(22):6603–6613.

48. Tarus B, Nguyen PH, Berthoumieu O, Faller P, Doig AJ, Derreumaux P. Molecular structure of the NQTrp inhibitor with the Alzheimer Aβ1-28 monomer. Eur J Med Chem. 2015 91:43–50.

49. Heller GT, Aprile FA, Michaels TCT, Limbocker R, Perni M, Ruggeri FS, Mannini B, Löhr T, Bonomi M, Camilloni C, De Simone A, Felli IC, Pierattelli R, Knowles TPJ, Dobson CM, Vendruscolo M. Small-molecule sequestration of amyloid-β as a drug discovery strategy for Alzheimer’s disease. Sci Adv. 2020 6(45):eabb5924.

50. Heller GT, Aprile FA, Bonomi M, Camilloni C, De Simone A, Vendruscolo M. Sequence specificity in the entropy-driven binding of a small molecule and a disordered peptide. J Mol Biol. 2017 429(18):2772–2779.

51. Herrera-Nieto P, Pérez A, De Fabritiis G. Small molecule modulation of intrinsically disordered proteins using molecular dynamics simulations. J Chem Inf Model. 2020 60(10):5003–5010.

52. Liang C, Savinov SN, Fejzo J, Eyles SJ, Chen J. Modulation of Amyloid-β42 Conformation by small molecules through nonspecific binding. J Chem Theory Comput. 2019 15(10):5169–5174.

53. Michel J, Cuchillo R. The impact of small molecule binding on the energy landscape of the intrinsically disordered protein C-myc. PLoS One. 2012 7(7):e41070.

54. Wang J, Wang W, Kollman PA, Case DA. Automatic atom type and bond type perception in molecular mechanical calculations. J Mol Graph Model. 2006 25(2):247–260.

55. Wang J, Wolf RM, Caldwell JW, Kollman PA, Case DA. Development and testing of a general amber force field. J Comput Chem. 2004 25(9):1157–1174.

56. Chelli R, Gervasio FL, Procacci P, Schettino V. Stacking and T-shape competition in aromatic-aromatic amino acid interactions. J Am Chem Soc. 2002 124(21):6133–6143.

57. Lamberto GR, Binolfi A, Orcellet ML, Bertoncini CW, Zweckstetter M, Griesinger C, Fernández CO. Structural and mechanistic basis behind the inhibitory interaction of PcTS on alpha-synuclein amyloid fibril formation. Proc Natl Acad Sci U S A. 2009 106(50):21057–21062.

58. Fonseca-Ornelas L, Eisbach SE, Paulat M, Giller K, Fernández CO, Outeiro TF, Becker S, Zweckstetter M. Small molecule-mediated stabilization of vesicle-associated helical α-synuclein inhibits pathogenic misfolding and aggregation. Nat Commun. 2014 5:5857.

59. Al-Hilaly YK, Biasetti L, Blakeman BJ, Pollack SJ, Zibaee S, Abdul-Sada A, Thorpe JR, Xue WF, Serpell LC. The involvement of dityrosine crosslinking in α-synuclein assembly and deposition in Lewy Bodies in Parkinson’s disease. Sci Rep. 2016 6:39171.

60. Ban D, Iconaru LI, Ramanathan A, Zuo J, Kriwacki RW. A small molecule causes a population shift in the conformational landscape of an intrinsically disordered protein. J Am Chem Soc. 2017 139(39):13692–13700.

