## Supporting Information for "Molecular basis of small-molecule binding to α-synuclein"

### Molecular basis of small-molecule binding to α-synuclein
### Supporting Information

#### Simulation Methods

Systems were equilibrated with GPU/Desmond.[1] Production runs at 300 K were performed in the NPT ensemble with Anton, a special-purpose machine capable of running very long MD simulations.[2] To further increase computational efficiency, we modified all the hydrogen (H) and water oxygen ($O_w$) masses (H: 4 Da; $O_w$: 10 Da); this allowed us to integrate the equations of motion using a RESPA scheme[3] with an inner time step of 4.5 fs and an outer time step of 9 fs.[4] Previous simulations of a well-characterized protein, villin, showed that using a larger time step and altering the masses of hydrogen and water oxygens did not have any substantial effect on the protein's kinetics or the thermodynamics.[5] Bonds involving hydrogen atoms were restrained to their equilibrium lengths using the M-SHAKE algorithm.[6] Nonbonded interactions were truncated at 10 Å, and long-range electrostatic interactions were computed using the *u*-series approach.[7] Proteins, water molecules, and ions were parameterized with the a99SB-*disp* force field[8] and all ligands were parameterized using the generalized Amber force field (GAFF).[9]

To calculate the dissociation constant ($K_D$) we used the expression[10]

$$K_{\mathrm{D}} = \frac{P_{\mathrm{u}}}{P_{\mathrm{b}}} v c^{\mathrm{o}} N_{\mathrm{Av}},$$

where $P_u$ is the fraction of simulation time in which the protein and ligand are unbound, $P_b$ is the fraction of simulation time in which the protein and ligand are bound, $v$ is the volume of the simulation box, $c^o$ is a standard-state concentration (1 mol $L^{-1}$), and $N_{Av}$ is Avogadro's number. To partition trajectories into bound and unbound states, we calculated the closest distance between the protein atoms and the ligand heavy atoms, and simulation frames in which this distance was less than 6 Å were considered bound.

We define aromatic stacking by considering the mutual arrangement of two aromatic groups as defined by a vector **R** connecting the centroids of the aromatic groups, and the angles α and β, defined by the normal of each aromatic plane and the vector **R** (Figure 2C). A conformation was classified as aromatic stacking if the length of **R** was <5 Å, α was >135 ° or <45 °, and β was >135° or <45°.[11]

Potential hydrogen donors were defined as all nitrogen, oxygen, or sulfur atoms with an attached hydrogen. Hydrogen bonds were identified with a distance cutoff of 3.5 Å and an angle <150°. Charge contacts were defined as being when any two atoms that contain opposite formal charges were within 5 Å. Hydrophobic contacts were defined as being when carbon atoms were within 4 Å (excluding backbone Cα atoms).

Mutual information (MI) was calculated using binary contact probabilities between all intermolecular interactions according to Shannon and Weaver[12]

$$\mathrm{I}(X;Y) = \sum_{y\in Y} \sum_{x\in X} p_{(X,Y)}\,(x,y) \log\left(\frac{p_{(X,Y)}(x,y)}{p_X(x)\,p_Y(y)}\right),$$

where $p_{(X,Y)}$ is the joint probability mass function of $X$ and $Y$; where $p_{(X)}$ and $p_{(Y)}$ are the marginal probability mass functions of $X$ and $Y$ respectively; and where $X$ and $Y$ represent binary contact probabilities of intermolecular interactions. Because the MI is strictly positive and bounded from above by the entropy of the joint distribution, the MI between low-entropy contact pairs will necessarily appear to be low even if they are perfectly coupled. To better resolve these contact pairs, we normalized the MI to the joint entropy of the contact pair,[13] which bounds the MI between 0 and 1.

#### Experimental Methods

The α-synuclein protein was expressed at 37 °C in *Escherichia coli* strain BL21(DE3) in M9 minimal medium supplemented with $^{15}NH_4Cl$ (Cambridge Isotope Laboratories), by induction with 1 mM IPTG. The cell pellet from 1 L cell culture was resuspended in 40 mL lysis buffer (10 mM Tris-HCl, pH 8, 1 mM EDTA, 0.5 M PMSF). Cell lysis was performed by three freeze/thaw cycles followed by eight ultrasonication cycles (20 sec at 90% power) on ice. The cell lysate was incubated at 96 °C in a water bath for 15 min. The supernatant was collected by centrifugation (Beckman Coulter, JA-25.5 rotor, 48,000 g at 4 °C for 45 min). Streptomycin sulfate was added to the supernatant at a final concentration of 10 mg $ml^{-1}$ and incubated at 4 °C for 30 min. The supernatant was collected by centrifugation (JA-25.5, 45 min, 48,000 g) and ammonium sulfate was added to a final concentration of 360 mg $mL^{-1}$ while stirring on ice for 30 min. After a final centrifugation step, the protein pellet was obtained and dialyzed against 25 mM Tris-HCl, pH 7.7 overnight. The dialysate was applied to a 30 mL POROS 20 HQ anion exchange column (Thermo Scientific) and eluted using a salt gradient from 0 to 1 M NaCl in the same buffer. In a final step the protein was further purified by size-exclusion chromatography on a HiLoad 16/60 Superdex 75 prep grade column (GE Healthcare) equilibrated with 50 mM HEPES, 100 mM NaCl, pH 7.4, 0.02% $NaN_3$ using an ÄKTA Purifier system (GE Healthcare). The final stock at 300 µM was filtered through sterile 0.22-µm filters and stored at −80 °C.

One-dimensional (1D) 1H NMR experiments and two-dimensional (2D) 1H-15N heteronuclear single quantum coherence (HSQC) experiments of α-synuclein in absence and presence of different concentrations of ligands were acquired at 288 K on Bruker 700 MHz and 800 MHz spectrometers equipped with triple-resonance 5 mm cryogenic probes. The protein concentration was 30 μM and all samples were in HEPES 50 mM pH 7.4, NaCl 100 mM, $D_2O$ 5%, $NaN_3$ 0.01% and DSS 50 μM. 1D experiments were used to calibrate different parameters. The signals of HEPES were always checked to correct pH changes caused by the addition of ligand powder through the titration. Signals from aromatic protons were used to measure the real concentration of each ligand. DSS methyl signal was used to calibrate the chemical shift. Spectra were processed with TopSpin 3.6 (Bruker) and analyzed using Sparky.[14] The CSP error is based on the resolution of the spectra.

Ligands 2, 5, 23, 47, and fasudil were purchased from commercial sources and used without additional purification. The (1D) 1H NMR characterization of these ligands is shown in Figure S12.

#### Supplementary References


1. Bergdorf, M., Baxter, S., Rendleman, C. A. & Shaw, D. E. Desmond/GPU performance as of November 2016. D. E. Shaw Research Technical Report DESRES/TR--2016-01 (2016)

2. David E. Shaw, J.P. Grossman, Joseph A. Bank, Brannon Batson, J. Adam Butts, Jack C. Chao, Martin M. Deneroff, Ron O. Dror, Amos Even, Christopher H. Fenton, Anthony Forte, Joseph Gagliardo, Gennette Gill, Brian Greskamp, C. Richard Ho, Douglas J. Ierardi, Lev Iserovich, Jeffrey S. Kuskin, Richard H. Larson, Timothy Layman, Li-Siang Lee, Adam K. Lerer, Chester Li, Daniel Killebrew, Kenneth M. Mackenzie, Shark Yeuk-Hai Mok, Mark A. Moraes, Rolf Mueller, Lawrence J. Nociolo, Jon L. Peticolas, Terry Quan, Daniel Ramot, John K. Salmon, Daniele P. Scarpazza, U. Ben Schafer, Naseer Siddique, Christopher W. Snyder, Jochen Spengler, Ping Tak Peter Tang, Michael Theobald, Horia Toma, Brian Towles, Benjamin Vitale, Stanley C. Wang, and Cliff Young, "Anton 2: Raising the Bar for Performance and Programmability in a Special-Purpose Molecular Dynamics Supercomputer," *Proceedings of the International Conference for High Performance Computing, Networking, Storage and Analysis (SC14)*, Piscataway, NJ: IEEE, 2014, pp. 41–53.

3. Tuckerman, M., Berne, B. J. & Martyna, G. J. Reversible multiple time scale molecular dynamics. *J. Chem. Phys.* **97(3),** 1990–2001 (1992)

4. Feenstra, K. A., Hess, B. & Berendsen, H. J. C. Improving efficiency of large time-scale molecular dynamics simulations of hydrogen-rich systems. *J. Comput. Chem.* **20(8),** 786–798 (1999)

5. Piana, S., Lindorff-Larsen, K. & Shaw, D. E. Atomic-level description of ubiquitin folding. *Proc. Natl. Acad. Sci. U. S. A.* **110(15),** 5915–5920 (2013).

6. Kräutler, V., van Gunsteren, W. F. & Hünenberger, P. H. A fast SHAKE algorithm to solve distance constraint equations for small molecules in molecular dynamics simulations. *J. Comput. Chem.* **22**, 501–508 (2001).

7. C. Predescu, A. K. Lerer, R. A. Lippert, B. Towles, J. P. Grossman, R. M. Dirks, D. E. Shaw, The *u* -series: A separable decomposition for electrostatics computation with improved accuracy. *J. Chem. Phys.* **152**, 084113 (2020).

8. Robustelli, P., Piana, S., and Shaw, D.E. Developing a molecular dynamics force field for both folded and disordered protein states. Proc Natl Acad Sci U S A. 2018 May 22;115(21):E4758-E4766.

9. Wang J, Wolf RM, Caldwell JW, Kollman PA, Case DA. Development and testing of a general amber force field. Journal of computational chemistry. 2004 Jul 15;25(9):1157-74

10. De Jong, D.H., Schäfer, L.V., De Vries, A.H., Marrink, S.J., Berendsen, H.J.C. and Grubmüller, H. (2011), Determining equilibrium constants for dimerization reactions from molecular dynamics simulations. J. Comput. Chem., 32: 1919-1928.

11. Chelli, R., Gervasio, F.L., Procacci, P. and Schettino, V., 2002. Stacking and T-shape competition in aromatic− aromatic amino acid interactions. *Journal of the American Chemical Society*, *124*(21), pp.6133-6143.

12. Shannon, C.E. and Weaver, W. (1949). The mathematical theory of communication. University of Illinois Press, Urbana, Illinois.

13. Malvestuto, F.M., 1986. Statistical treatment of the information content of a database. *Information Systems*, *11*(3), pp.211-223

14. Lee, W., Tonelli, M. & Markley, J.L. NMRFAM-SPARKY: enhanced software for biomolecular NMR spectroscopy. Bioinformatics 31, 1325-1327 (2015).

15. LeVine MV, Weinstein H (2014) NbIT - A New Information Theory-Based Analysis of Allosteric Mechanisms Reveals Residues that Underlie Function in the Leucine Transporter LeuT. PLOS Computational Biology 10(5): e1003603.

#### Supplementary Tables

| Side chain | Quadrant | | | | Max/Min |
|---|---|---|---|---|---|
| | 1 | 2 | 3 | 4 | |
| **Y125** | 0.254 | 0.232 | 0.271 | 0.243 | 1.2 |
| **Y125 with D135 contact** | 0.441 | 0.273 | 0.178 | 0.108 | 4.1 |
| **Y133** | 0.254 | 0.191 | 0.267 | 0.287 | 1.5 |
| **Y133 with D135 contact** | 0.156 | 0.131 | 0.127 | 0.586 | 4.6 |
| **Y136** | 0.206 | 0.234 | 0.277 | 0.283 | 1.4 |
| **Y136 with D135 contact** | 0.248 | 0.357 | 0.235 | 0.160 | 2.2 |

**fasudil**

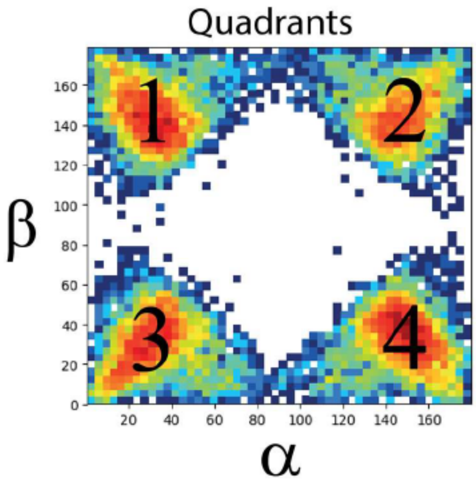


| Side chain | Quadrant | | | | Max/Min |
|---|---|---|---|---|---|
| | 1 | 2 | 3 | 4 | |
| **Y125** | 0.068 | 0.252 | 0.280 | 0.400 | 5.8 |
| **Y125 with D135 contact** | 0.190 | 0.312 | 0.184 | 0.314 | 1.7 |
| **Y133** | 0.270 | 0.230 | 0.268 | 0.232 | 1.2 |
| **Y133 with D135 contact** | 0.263 | 0.255 | 0.213 | 0.269 | 1.3 |
| **Y136** | 0.174 | 0.183 | 0.292 | 0.350 | 2.0 |
| **Y136 with D135 contact** | 0.044 | 0.040 | 0.386 | 0.530 | 13.2 |

**Ligand 47**

**Table S1**. Protein-ligand aromatic stacking orientations. For all conformations where the ring centers of a ligand and a tyrosine phenol group were within 5 Å, we calculated the angles ($\alpha$,$\beta$) of the normal vectors of each ring plane relative to a vector connecting the two ring centers (illustrated in Figure 2C). To quantify asymmetries in the distributions of these angles, we report the ratio of the populations of the most populated quadrant to the least populated quadrant.

| Ligand | $K_D$ full length (mM) | $K_D$ 29–49 (mM) | $K_D$ 121–140 (mM) |
|---|---|---|---|
| **fasudil** | 2.98 +/- 0.022 | 15.9 +/- 0.15 | 6.80 +/- 0.080 |
| **47** | 2.26 +/- 0.039 | 10.3 +/- 0.18 | 5.0 +/- 0.12 |
| **23** | 6.4 +/- 0.13 | 24 +/- 1.2 | 25.5 +/- 0.98 |

**Table S2.** Simulated $K_D$ values of fasudil, Ligand 47, and Ligand 23 to full-length α-synuclein, and two subregions containing tyrosine residues.

| Ligand | SMILES string of compound | $K_D$ α-syn 29–49 (mM) | Blocking Error | $K_D$ α-syn-C-term (mM) | Blocking Error |
|---|---|---|---|---|---|
| 1 | S(=O)(=O)(N1CCNCCC1)c1c2c(ccc1)cccc2 | 35.327 | 1.649 | 8.625 | 0.332 |
| 2 | S(=O)(=O)(N1CCNCCC1)c1cc(C(=O)NC(C)c2ccccc2)ccc1 | 31.311 | 1.229 | 7.193 | 0.281 |
| 3 | S(=O)(=O)(N1CCNCCC1)c1cc2c(cc1)cccc2 | 28.126 | 1.165 | 6.107 | 0.366 |
| 4 | Clc1c(Cl)ccc(S(=O)(=O)N2CCNCCC2)c1 | 34.908 | 1.106 | 7.96 | 0.257 |
| 5 | Clc1c(Cl)cccc1S(=O)(=O)N1CCNCCC1 | 36.165 | 1.346 | 7.788 | 0.265 |
| 6 | S(=O)(=O)(N1CCNCCC1)c1ccc(C2CCCCC2)cc1 | 31.151 | 1.311 | 7.32 | 0.265 |
| 7 | S(=O)(=O)(N1CCNCCC1)c1c2ncc(C)cc2ccc1 | 38.709 | 1.462 | 8.281 | 0.276 |
| 8 | S(=O)(=O)(N1CCNCCC1)c1c2c(c(OC)cc1)cccc2 | 32.208 | 1.317 | 6.99 | 0.453 |
| 9 | S(=O)(=O)(N1CCNCCC1)c1ccc(Oc2ccc(OC)cc2)cc1 | 23.828 | 1.139 | 6.018 | 0.31 |
| 10 | Clc1cc2c(S(=O)(=O)N3CCNCCC3)csc2cc1 | 28.589 | 1.295 | 6.649 | 0.376 |
| 11 | S(=O)(=O)(N1CCNCCC1)c1c2c(c(C)cc1)nccc2 | 27.541 | 1.249 | 6.321 | 0.343 |
| 12 | S(=O)(=O)(N1CCNCCC1)c1cc2sc(C(C)(C)C)nc2cc1 | 28.25 | 1.22 | 5.035 | 0.253 |
| 13 | Clc1c(Oc2ccc(S(=O)(=O)N3CCNCCC3)cc2)cccc1 | 25.053 | 1.57 | 5.459 | 0.21 |
| 14 | S(=O)(=O)(N1CCNCCC1)c1cc2C(=O)NCCc2cc1 | 37.108 | 1.889 | 7.225 | 0.243 |
| 15 | Clc1cc2c(sc(S(=O)(=O)N3CCNCCC3)c2)cc1 | 28.361 | 1.646 | 5.476 | 0.324 |
| 16 | S(=O)(=O)(N1CCNCCC1)c1ccc(Oc2cc(C)ccc2)cc1 | 27.766 | 1.292 | 6.296 | 0.354 |
| 17 | S(=O)(=O)(N1CCNCCC1)c1cn(-c2ccccc2)nc1 | 34.5 | 1.553 | 7.074 | 0.267 |
| 18 | S(=O)(=O)(N1CCNCCC1)c1c2c(c(C(=O)OC)ncc2)ccc1 | 27.247 | 1.341 | 4.522 | 0.287 |
| 19 | S(=O)(=O)(N1CCNCCC1)c1c(C)n(C)c2c1cccc2 | 35.938 | 1.404 | 8.78 | 0.33 |
| 20 | S(=O)(=O)(N1CCNCCC1)c1cc(-c2scc(C)n2)ccc1 | 28.602 | 1.22 | 5.519 | 0.297 |
| 21 | S(=O)(=O)(N1CCN(C)CCC1)c1c2c(cncc2)ccc1 | 37.696 | 1.96 | 8.527 | 0.374 |
| 22 | S(=O)(=O)(N1CC(N)CC1)c1c2c(cncc2)ccc1 | 35.593 | 1.574 | 7.791 | 0.286 |
| 23 | S(=O)(=O)(N1CCC(O)CC1)c1c2c(cncc2)ccc1 | 24.399 | 0.985 | 18.402 | 0.576 |

| | | | | | |
|---|---|---|---|---|---|
| 24 | S(=O)(=O)(N1CCC(CN)CC1)c1c2c(cncc2)ccc1 | 28.919 | 1.216 | 6.575 | 0.241 |
| 25 | S(=O)(=O)(N1CCC(N)CC1)c1c2c(cncc2)ccc1 | 33.855 | 1.53 | 7.578 | 0.376 |
| 26 | S(=O)(=O)(N1CC(CN)CCC1)c1c2c(cncc2)ccc1 | 29.836 | 1.231 | 7.015 | 0.309 |
| 27 | S(=O)(=O)(N1CC(CN)CC1)c1c2c(cncc2)ccc1 | 34.5 | 1.394 | 6.935 | 0.296 |
| 28 | S(=O)(=O)(N1CCNCC1)c1c2c(cncc2)ccc1 | 35.829 | 1.423 | 6.68 | 0.441 |
| 29 | S(=O)(=O)(N1CC(N)CCC1)c1c2c(cncc2)ccc1 | 34.08 | 1.823 | 8.382 | 0.341 |
| 30 | S(=O)(=O)(N1CCOCC1)c1c2c(cncc2)ccc1 | 25.787 | 0.999 | 18.462 | 0.719 |
| 33 | S(=O)(=O)(N1CCNCCC1)c1ccc(-c2ccccc2)cc1 | 27.261 | 1 | 6.181 | 0.283 |
| 34 | S(=O)(=O)(N1CCNCCC1)c1c2c(nccc2)ccc1 | 36.043 | 1.891 | 8.549 | 0.287 |
| 35 | S(=O)(=O)(N1CCNCCC1)c1c2ncccc2ccc1 | 40.707 | 1.944 | 9.64 | 0.368 |
| 36 | S(=O)(=O)(N1CCNCCC1)c1cc(-c2ccccc2)ccc1 | 27.212 | 1.368 | 6.111 | 0.456 |
| 37 | S(=O)(=O)(N1CCNCCC1)c1c(-c2ccccc2)cccc1 | 41.773 | 2.18 | 10.448 | 0.321 |
| 38 | S(=O)(=O)(N1CCNCCC1)c1ccccc1 | 48.861 | 1.838 | 11.681 | 0.332 |
| 39 | S(=O)(=O)(N1CCNCCC1)c1c2c([nH]c1)nccc2 | 37.085 | 1.691 | 7.592 | 0.408 |
| 40 | S(=O)(=O)(N1CCNCCC1)c1ccc(C(=O)Nc2ccccc2)cc1 | 24.884 | 0.981 | 6.003 | 0.333 |
| 41 | S(=O)(=O)(N1CCNCCC1)c1c2c(cncc2)ccc1 | 36.892 | 1.372 | 7.866 | 0.155 |
| 46 | S(=O)(=O)(N1CCNCCC1)c1c2nsnc2ccc1 | 39.878 | 1.627 | 8.91 | 0.264 |
| 47 | S(=O)(=O)(N1CCNCCC1)c1cc2c(N(C(=O)C)CC2)cc1 | 28.849 | 1.736 | 4.51 | 0.166 |
| 48 | S(=O)(=O)(N1CCNCCC1)c1cc2OC(=O)N(C)c2cc1 | 30.368 | 1.44 | 5.701 | 0.277 |
| 49 | S(=O)(=O)(N1CCNCCC1)c1cc2N(C(=O)C)CCc2cc1 | 27.155 | 1.343 | 5.326 | 0.287 |
| 50 | S(=O)(=O)(N1CCNCCC1)c1cn(Cc2ccccc2)nc1 | 37.22 | 2.068 | 8.854 | 0.482 |
| 51 | S(=O)(=O)(Cc1c2ncccc2ccc1)N1CCNCCC1 | 36.109 | 2.252 | 8.027 | 0.341 |
| 52 | S(=O)(=O)(N1CCNCCC1)c1cnc2n(C)nc(C)c2c1 | 32.342 | 1.34 | 5.305 | 0.286 |
| 53 | S(=O)(=O)(N1CCNCCC1)c1ccc(OCc2n(C)ncn2)cc1 | 30.392 | 1.36 | 7.116 | 0.302 |
| 54 | S(=O)(=O)(N1CCNCCC1)c1cnc2onc(CC)c2c1 | 38.974 | 1.451 | 8.09 | 0.3 |
| 55 | S(=O)(=O)(N1CCNCCC1)c1c2nn(C)nc2ccc1 | 45.645 | 1.749 | 10.638 | 0.342 |

| | | | | | |
|---|---|---|---|---|---|
| 56 | S(=O)(=O)(N1CCNCCC1)c1cc2NC(=O)Oc2cc1 | 35.788 | 2.274 | 6.182 | 0.317 |

**Table S3.** $K_D$ values of all small-molecule compounds simulated in this work with fragments of α-syn containing either residues 29–49 or residues 121–140 (α-syn-C-term). All simulations were 60 μs in length, except for those of fasudil and Ligand 47 with α-syn-C-term which were 200 μs in length.

#### Supplementary Figures

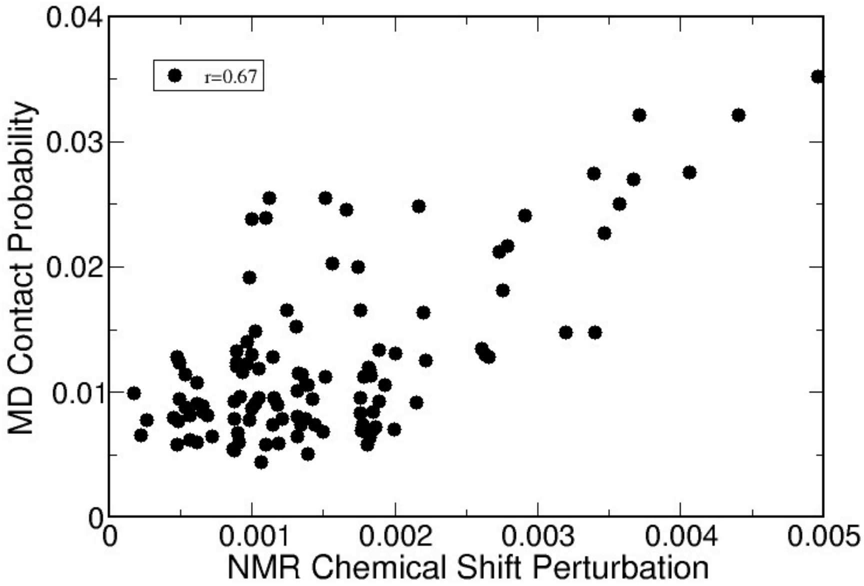


**Figure S1**. Correlation of the contact probability between each residue of α-synuclein and fasudil observed in an unbiased 1.5-ms MD simulation (Fig. 1A) and NMR chemical shift perturbations observed in the presence of 2.7 mM fasudil (Fig. 1B).

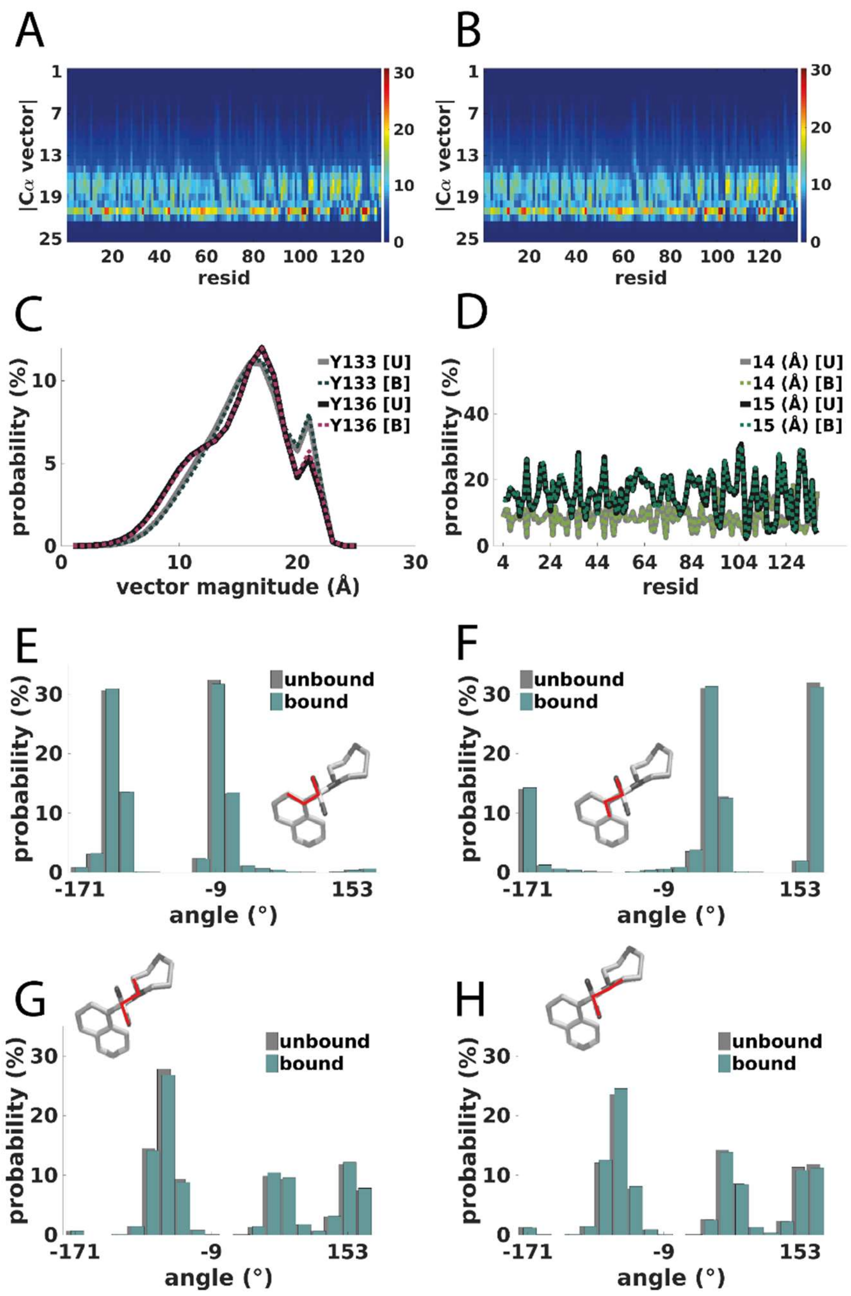


**Figure S2.** No substantial difference exists between the conformational ensembles of bound and unbound states of fasudil and α-syn for either the ligand or the protein. Backbone conformation distributions, quantified by the magnitude of the cross product of the vectors between sequential backbone carbon alphas, |Cα vector| = $|(C\alpha_i - C\alpha_{i-1}) \times (C\alpha_i - C\alpha_{i+1})|$, are unchanged between bound (A) and unbound ensembles (B). The color bars in (A) and (B) indicate the percentage probability that the |Cα vector| is the given magnitude (*y*-axis) for each residue (*x*-axis). The bound and unbound distributions of Y133 Cα and Y136 Cα, are shown in (C), where solid lines represent the unbound distributions [U] and dotted lines represent the bound distributions [B]. The distributions of the unbound and bound distributions of |Cα vector| magnitudes at 14 $Å^2$ and 15 $Å^2$ are shown in (D). Finally, bound and unbound distributions of four central dihedrals of fasudil, highlighted in red in the ligand cartoons, are shown in (E) through (H).

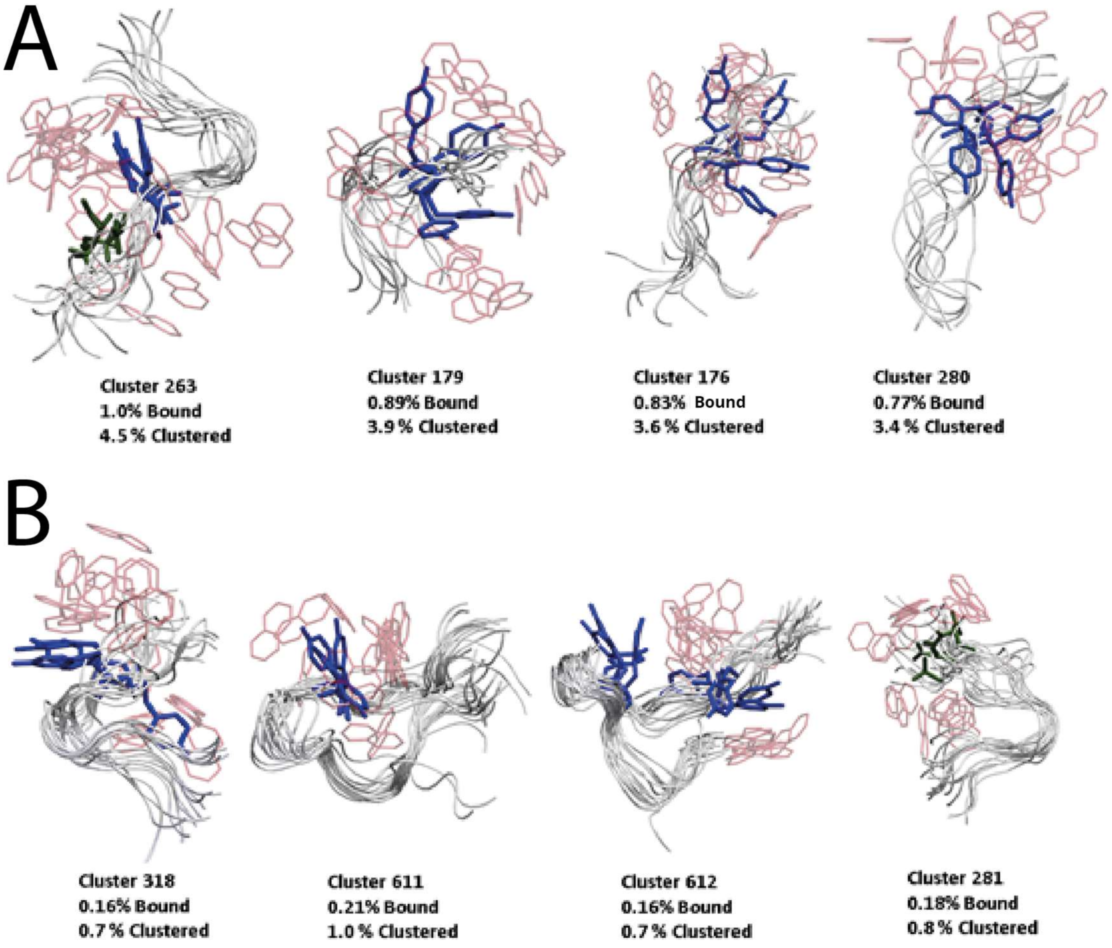


**Figure S3**. A clustering algorithm was applied to all frames containing interactions between fasudil and residues 121–140 of α-synuclein to find regions of local backbone ordering. The four largest clusters with 1 or 2 ordered residues (A) or at least 4 ordered residues (B) are depicted here. Ordered residues are those with |Cα vector|-associated stability within 180 ns. Backbone conformations with stable residues are shown as white tubes, and representative conformations of side chains with highest fraction of interactions with fasudil in each cluster are shown (tyrosine residues are colored blue, and D135 is colored green). Representative orientations of the isoquinoline ring of fasudil are shown in red, illustrating the highly heterogeneous ensemble of binding modes observed in each cluster. Binding modes are very heterogeneous even in the most stable α-syn conformations with at least four stable residues.

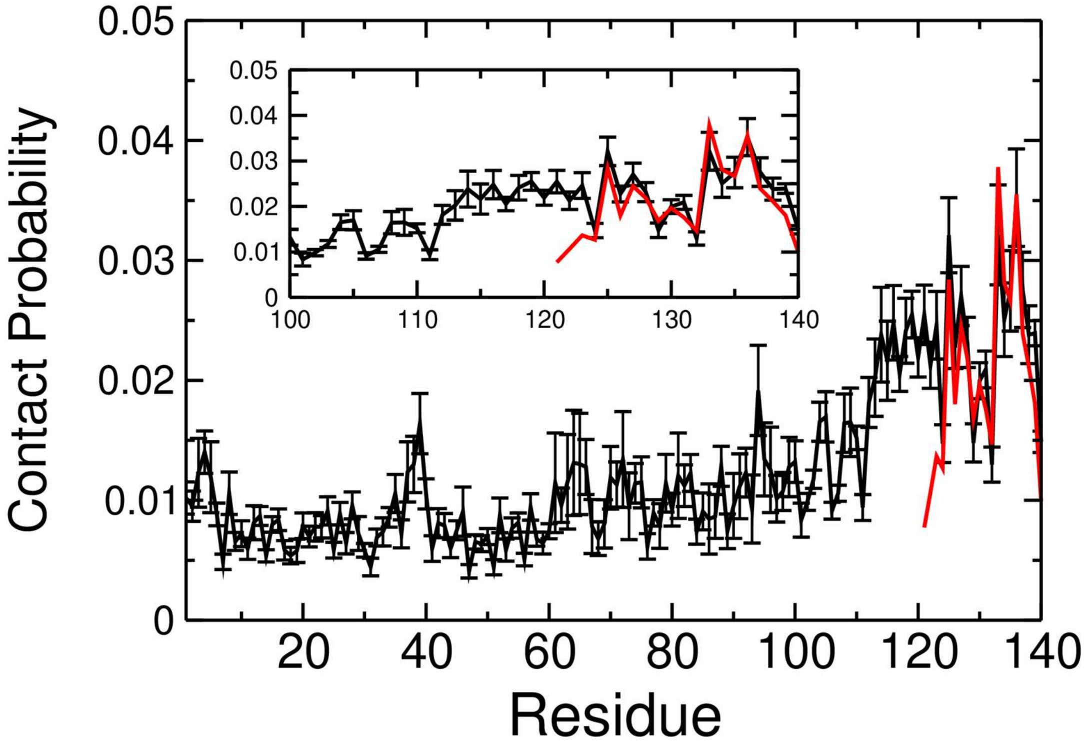


**Figure S4**. Simulations with the α-syn-C-term construct produced a similar contact probability with fasudil when compared to full-length α-syn. Contact probability for fasudil and truncated α-syn-C-term (red) is overlaid onto the full-length contact probability from Fig. 1a. The inset is zoomed into residues 100–140.

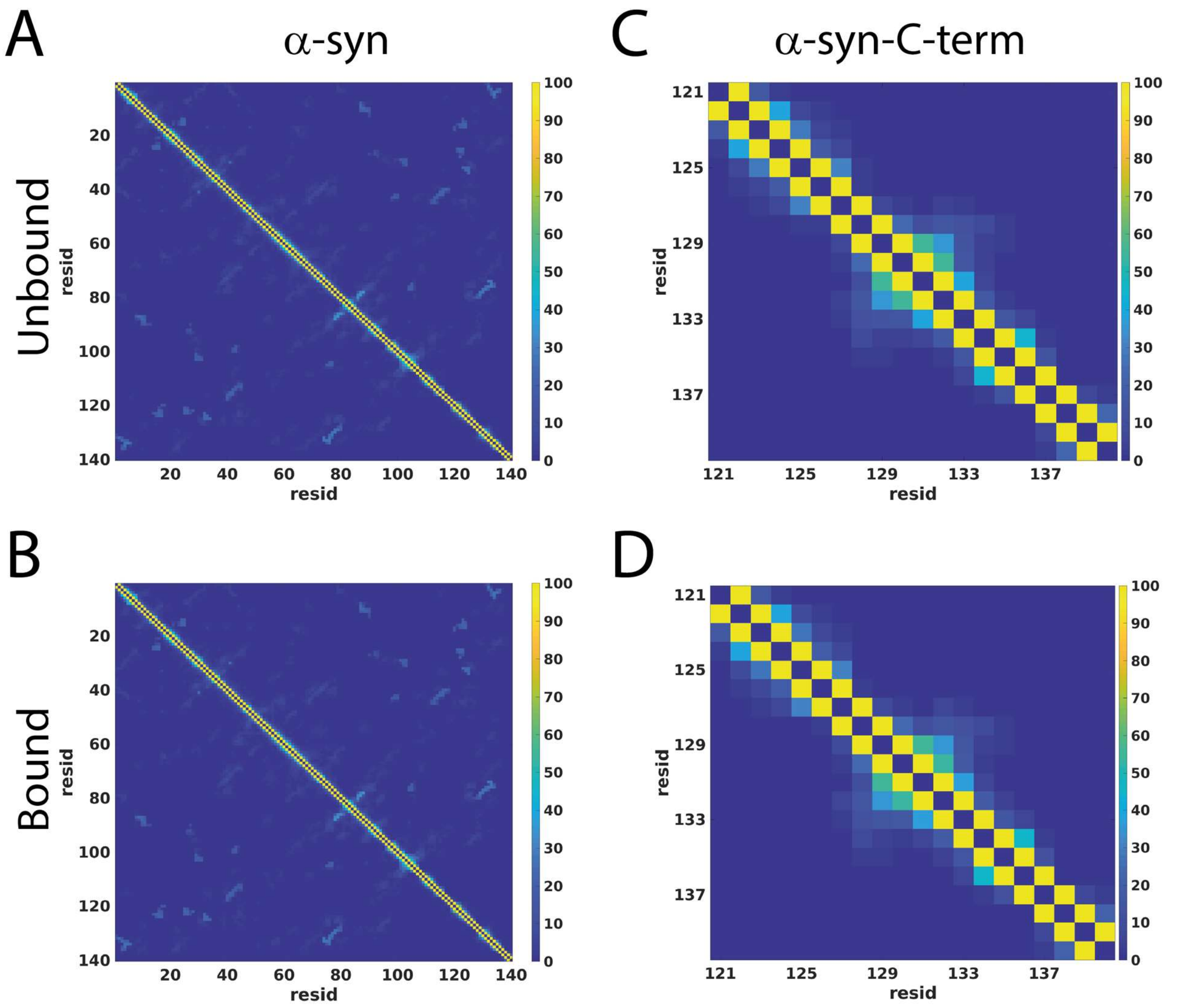


**Figure S5**. Protein-protein contact maps with fasudil unbound and bound for full length α-syn (A, B) and α-syn-C-term (C, D). Two protein residues are in contact if their Cα-Cα distance is less than 6 Å. The color bar denotes the percentage of simulations frames that contain the contact.

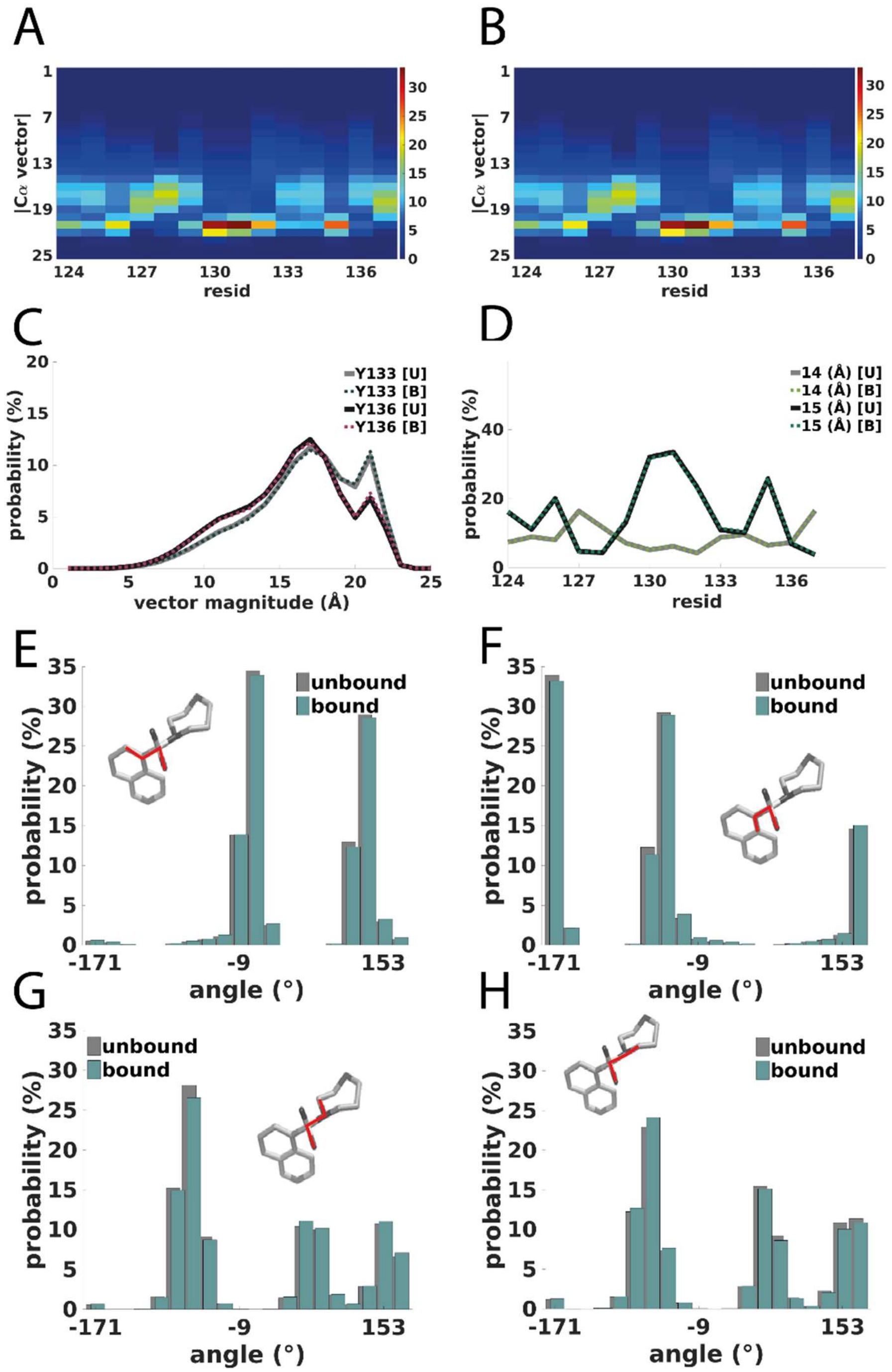


**Figure S6.** Same as Fig. S2, but for α-syn-C-term.

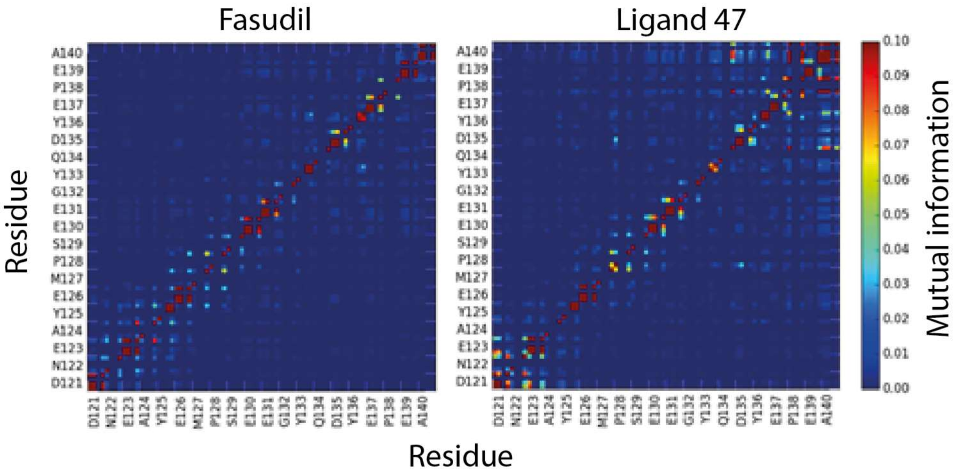


**Figure S7**. Comparison of mutual information for all interactions observed in the bound ensembles for fasudil (left) and Ligand 47 (right). For each residue, the MI between each potential ligand interaction (hydrophobic contacts, aromatic stacking, charge-charge, and hydrogen bonding) and all other potential ligand interactions is shown. We calculated the MI using binary contact probabilities according to ref. 15. Because the MI is strictly positive and bounded from above by the entropy of the joint distribution, the MI between low-entropy contact pairs will necessarily appear to be low even if they are perfectly coupled. To better resolve these contact pairs, we normalized the MI to the joint entropy of the contact pair, which bounds the MI between 0 and 1.

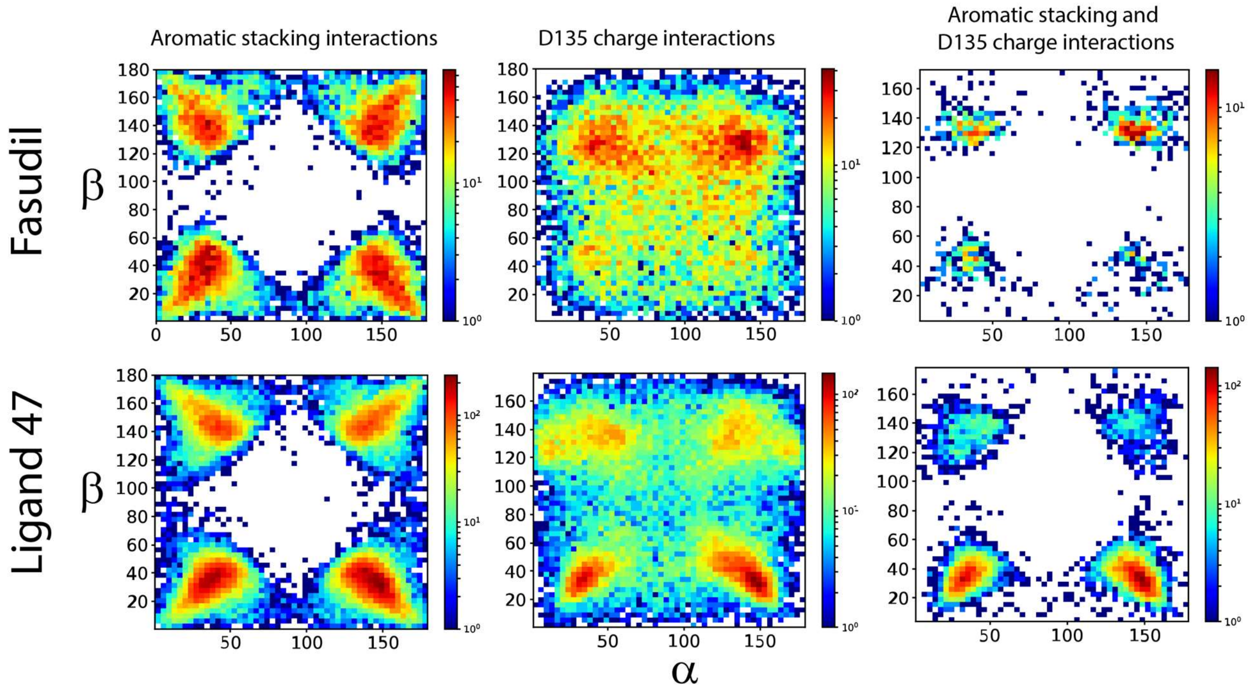


**Figure S8**. Ligand 47 orients stacking orientations more strongly than fasudil when charge contacts are made. Stacking orientation between the aromatic rings of fasudil (top row) and Ligand 47 (bottom row) with the side chain of Y136. The plots correspond to orientations of the ligand rings with Y136 when aromatic stacking interactions are formed (first column, see Fig. 2C), when D135 charge interactions are formed (second column), and when both aromatic stacking and D135 charge interactions are formed simultaneously (third column).

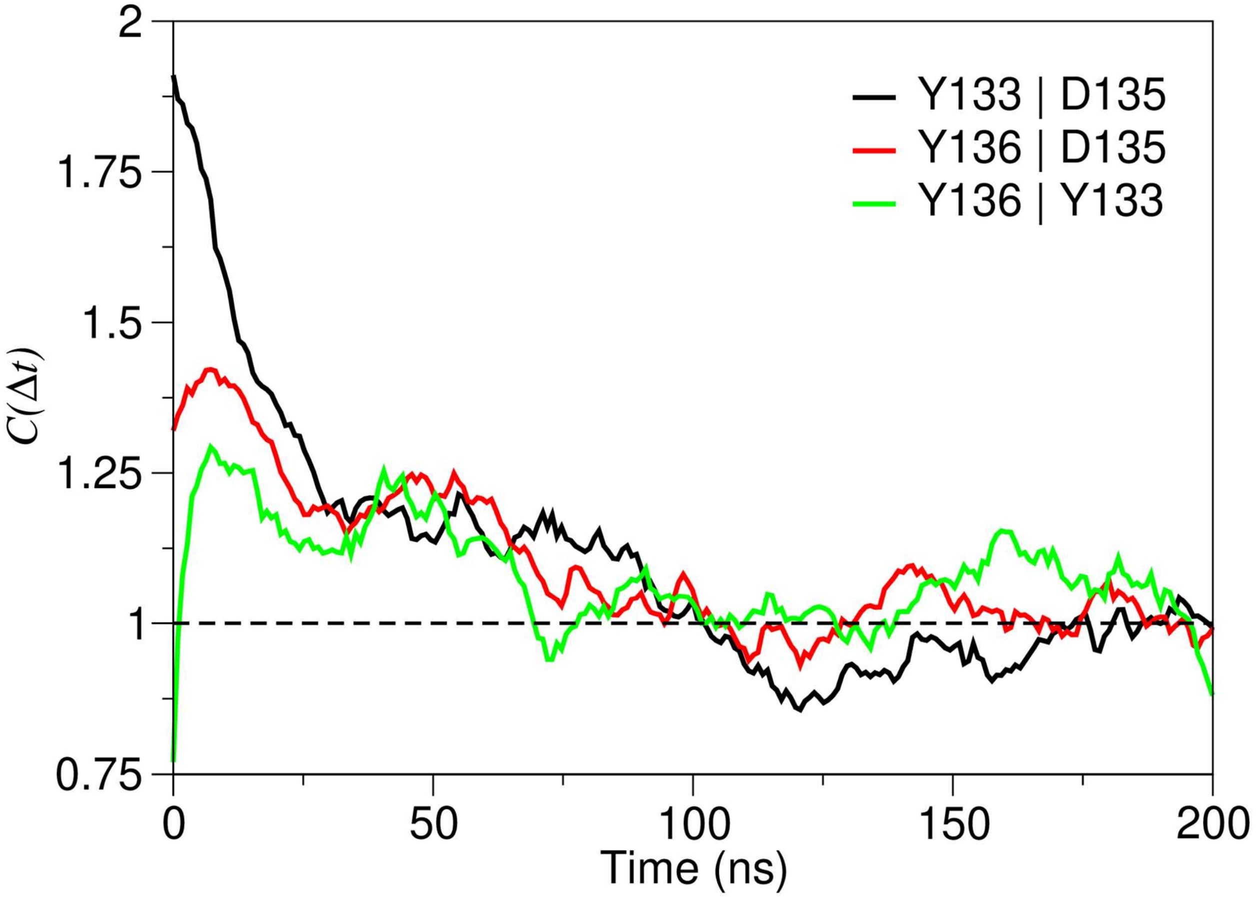


**Figure S8B**. Temporal correlations for fasudil forming π-stacking interactions with Y133 and Y136; charge-charge interactions with D135 persist for as long as 100 ns. The time correlation function, $C(\Delta t) = \frac{P(A; t_0 + \Delta t | B; t_0)}{P(A)}$, measures the probability of an interaction between fasudil and residue A at time $t_0 + \Delta t$, given there being an interaction between fasudil and residue B at time $t_0$. The normalization is such that as $\Delta t \to 0$, $C(\Delta t) \to \frac{P(A|B)}{P(A)} = \frac{P(A \cap B)}{P(A)P(B)}$, which is a measure of the correlation of the interactions with residues A and B (a value of 1 indicates that the probabilities of fasudil interacting with residue A and residue B are not correlated). As $\Delta t \to \infty$, $C(\Delta t) \to \frac{P(A)}{P(A)} = 1$, because, at long times, there is no longer any correlation between interactions with A and with B. The plot above shows $C(\Delta t)$ for the conditional probability of the interactions listed in the legend. Note that simultaneous interactions do not contribute significantly to these correlations, since the persistence of such interactions is only on the order of several hundred picoseconds: The average persistence times of simultaneous protein-ligand interactions with Y133 and D135, Y136 and D135, and Y133 and Y136, are 570 ps, 350 ps, and 300 ps, respectively. These calculations were performed with fasudil interacting with α-syn-C-term.

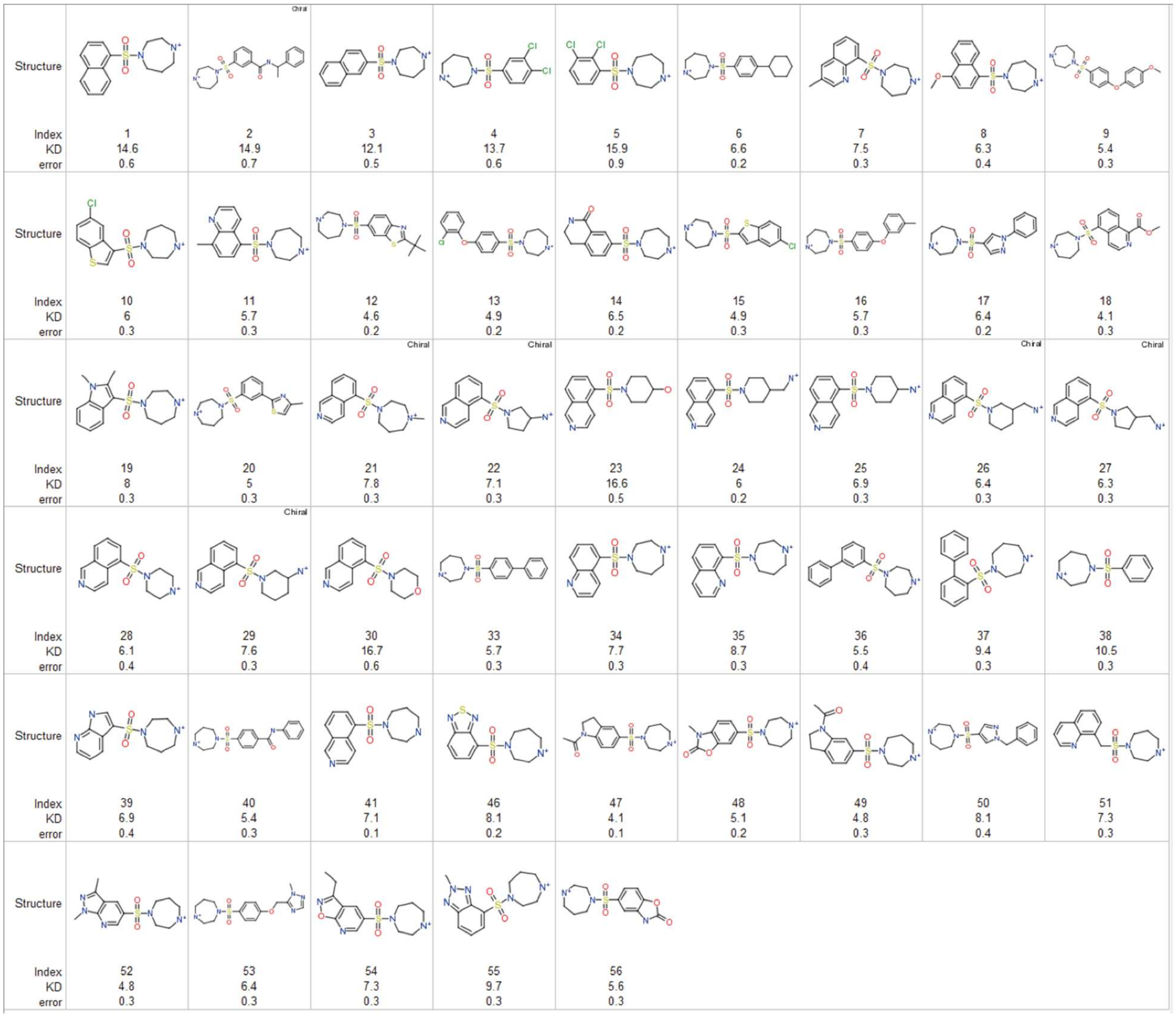

| | | | | | | | | | |
|---|---|---|---|---|---|---|---|---|---|
| Index | 1 | 2 | 3 | 4 | 5 | 6 | 7 | 8 | 9 |
| KD | 14.6 | 14.9 | 12.1 | 13.7 | 15.9 | 6.6 | 7.5 | 6.3 | 5.4 |
| error | 0.6 | 0.7 | 0.5 | 0.6 | 0.9 | 0.2 | 0.3 | 0.4 | 0.3 |
| Index | 10 | 11 | 12 | 13 | 14 | 15 | 16 | 17 | 18 |
| KD | 6 | 5.7 | 4.6 | 4.9 | 6.5 | 4.9 | 5.7 | 6.4 | 4.1 |
| error | 0.3 | 0.3 | 0.2 | 0.2 | 0.2 | 0.3 | 0.3 | 0.2 | 0.3 |
| Index | 19 | 20 | 21 | 22 | 23 | 24 | 25 | 26 | 27 |
| KD | 8 | 5 | 7.8 | 7.1 | 16.6 | 6 | 6.9 | 6.4 | 6.3 |
| error | 0.3 | 0.3 | 0.3 | 0.3 | 0.5 | 0.2 | 0.3 | 0.3 | 0.3 |
| Index | 28 | 29 | 30 | 33 | 34 | 35 | 36 | 37 | 38 |
| KD | 6.1 | 7.6 | 16.7 | 5.7 | 7.7 | 8.7 | 5.5 | 9.4 | 10.5 |
| error | 0.4 | 0.3 | 0.6 | 0.3 | 0.3 | 0.3 | 0.4 | 0.3 | 0.3 |
| Index | 39 | 40 | 41 | 46 | 47 | 48 | 49 | 50 | 51 |
| KD | 6.9 | 5.4 | 7.1 | 8.1 | 4.1 | 5.1 | 4.8 | 8.1 | 7.3 |
| error | 0.4 | 0.3 | 0.1 | 0.2 | 0.1 | 0.2 | 0.3 | 0.4 | 0.3 |
| Index | 52 | 53 | 54 | 55 | 56 | | | | |
| KD | 4.8 | 6.4 | 7.3 | 9.7 | 5.6 | | | | |
| error | 0.3 | 0.3 | 0.3 | 0.3 | 0.3 | | | | |

**Figure S9**. Compounds selected to probe how chemical modifications affected the affinity of the interaction and the simulated binding mechanisms. Each compound was simulated with a fragment of α-synuclein containing residues 121 to 140. The $K_D$ values calculated from these simulations are shown. Commercially available compounds were chosen; Ligand 41 is fasudil.

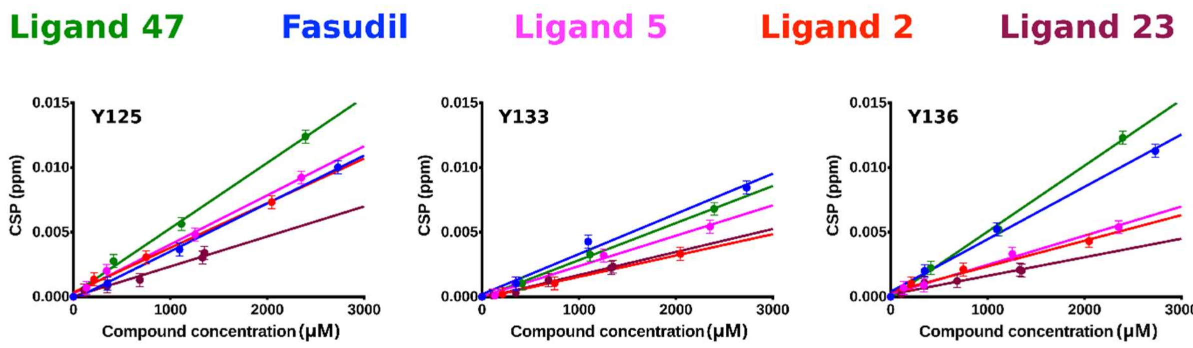


**Figure S10**. Individual CSP titration curves for Y125, Y133, and Y136, as a function of compound concentration for Ligand 47, fasudil, Ligand 5, Ligand 2, and Ligand 23.

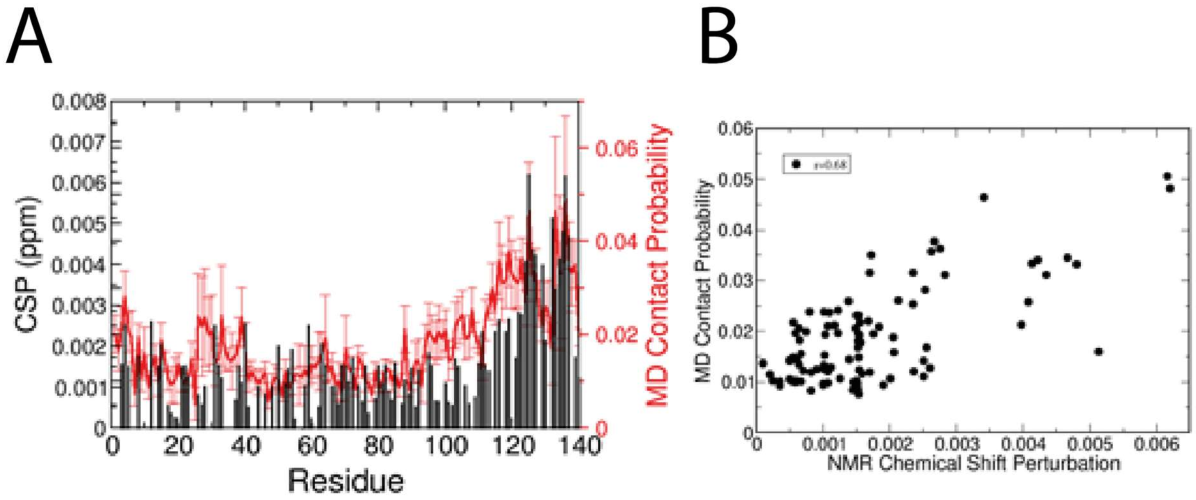


**Figure S11**. (A) Contact probabilities (red) computed from an unbiased MD simulation plotted against NMR chemical shift perturbations of Ligand 47 with α-synuclein. (B) Correlation between the contact probability of each residue of α-synuclein and Ligand 47 and NMR chemical shift perturbations observed in the presence of 2.4 mM Ligand 47.

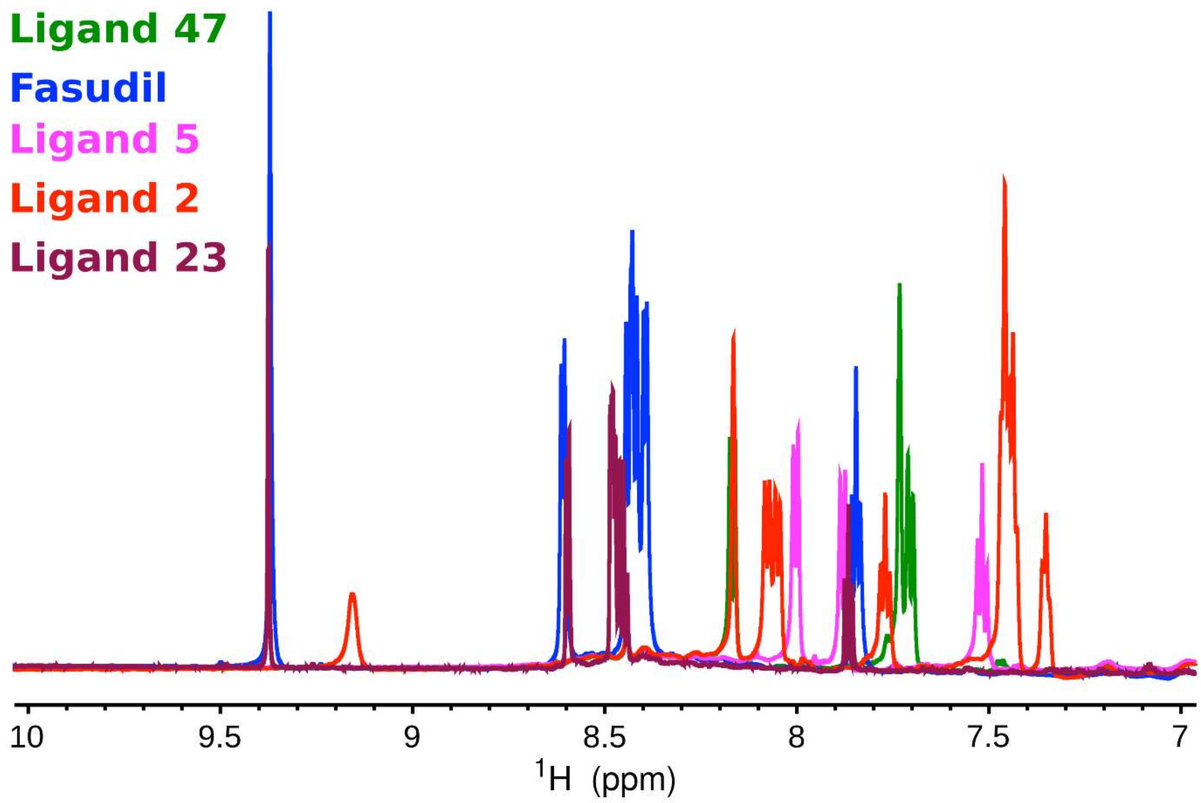


**Figure S12**. 1D NMR spectra of measured compounds. Spectra were acquired during titration experiments in the presence of α-synuclein, which had a low signal relative to the compounds, and HEPES buffer, which had a high signal relative to the compounds, so only the region of aromatics and amide nitrogens between 10 and 7 ppm is plotted, as HEPES has no signal there. The concentration of the compounds are the maximum used in the titration experiments (same as in the HSQC figures in Fig. 4). Note that the spectra of fasudil and Ligand 23 are very similar because their aromatic moiety is the same.
